# A human homolog of SIR2 antiphage proteins mediates immunity via the TLR pathway

**DOI:** 10.1101/2024.09.18.613514

**Authors:** Delphine Bonhomme, Hugo Vaysset, Eirene Marie Q. Ednacot, Vasco Rodrigues, Jean Cury, Veronica Hernandez Trejo, Philippe Benaroch, Benjamin R. Morehouse, Aude Bernheim, Enzo Z. Poirier

## Abstract

The full extent of immune system conservation between prokaryotes and eukaryotes is unknown. However, recent research supports that a subset of bacterial antiphage proteins is conserved in eukaryotes and likely gave rise to key actors of mammalian immunity. Here, we show that the SIR2 protein domain, present in bacterial antiphage systems, plays a role in eukaryotic innate immunity. Through phylogenetic analysis, we identify SIRanc, a human protein with a SIRim domain (subtype of SIR2). We demonstrate that SIRanc plays a pivotal role in the animal toll-like receptor (TLR) pathway of innate immunity by mediating the transcriptional upregulation of proinflammatory genes downstream of TLR stimulation. This depends on the enzymatic activity of SIRanc, which degrades NAD^+^, a central cellular metabolite. Finally, we show that proteins with a SIRim domain are diverse and widespread, detected in 19% of eukaryotic genomes, with SIRanc representing one of the five sirim lineages. This work opens avenues of research on the potential role of eukaryotic SIRim proteins in immunity, as well as on the involvement of SIRanc in human pathology.

## Introduction

The arms race between hosts and pathogens drives continuous diversification of immune mechanisms. As bacterial, archaeal and eukaryotic organisms face distinct threats, research thus far has largely focused on clade-specific immunity. However, recent work uncovered immune mechanisms shared between prokaryotes and eukaryotes, including humans, revealing an overlooked aspect of immunity: conservation (*1–3*). This conservation includes the cGAS-STING pathway, which is pivotal in the detection of DNA viruses, as well as caspases/gasdermins, which regulate cell death, and viperins, central antiviral effectors (*4–9*). Similarly, toll-interleukin-1 receptor (TIR) domains are involved in antiphage systems as well as in animal immune pathways (*10–12*). In prokaryotes and plants, TIR domains perform an enzymatic function, degrading NAD^+^ to trigger cell demise or to produce immune secondary messengers (*10*, *12–14*). In mammals, toll-like receptors (TLRs), which are crucial in detecting pathogen cues upon infection, transduce immune signals via a cytosolic TIR domain (*11*). The breadth of conservation of immune actors across domains of life remains to be investigated.

Prokaryotes and eukaryotes encode proteins bearing a Silent Information Regulator 2 (SIR2) domain, originally described in the sirtuin family of enzymes (*15*). By performing NAD-dependent deacetylation, sirtuins play an important role in regulating gene expression via the modification of histones in eukaryotes, or through the regulation of metabolism in bacteria (*15*, *16*). Recently, a novel function was uncovered for the SIR2 domain in bacteria: a role in antiphage defence (*12*, *17–20*). Multiple antiphage systems encode a SIR2 domain that provides immunity through degradation of cellular NAD^+^. For example, phage-mediated activation of the protein Thoeris A (ThsA) leads to SIR2-dependent depletion of NAD^+^ and growth arrest of the infected cell (*12*, *19*, *20*). SIR2 domain-containing proteins thus perform immune and non-immune functions in bacteria. Whether SIR2 domains from antiphage systems are conserved in eukaryotes and contribute to human immunity is currently unknown.

## Results

### Two human proteins are homologs of SIRim antiphage proteins

We set out to identify all prokaryotic and eukaryotic proteins containing a SIR2 domain encoded in publicly available sequenced genomes. We first implemented homology-based detection of SIR2 proteins that are unrelated to bacterial, archaeal, or eukaryotic immunity, and of SIR2 proteins involved in bacterial antiphage systems. To do so, we built a general protein profile for the SIR2 family using 4,302 SIR2 homologs from across the tree of life (see Methods) and used it to search a database of 30,254 bacterial, archaeal and eukaryotic genomes. This homology search identified 57,911 SIR2 proteins across kingdoms (29,634 in bacteria, 2,094 in archaea, and 26,183 in eukaryotes, Table S1). To better characterise the diversity within the SIR2 family, we constructed a phylogenetic tree (Fig 1a, see Methods). In this tree, SIR2 proteins are clustered into two monophyletic groups (Fig 1a, beige and brown), with each group composed of bacterial, archaeal, and eukaryotic proteins (inner circle, blue, grey and green, respectively). This indicates that the family of SIR2 proteins can be partitioned into two subfamilies of ancient origin.

**Figure 1.**
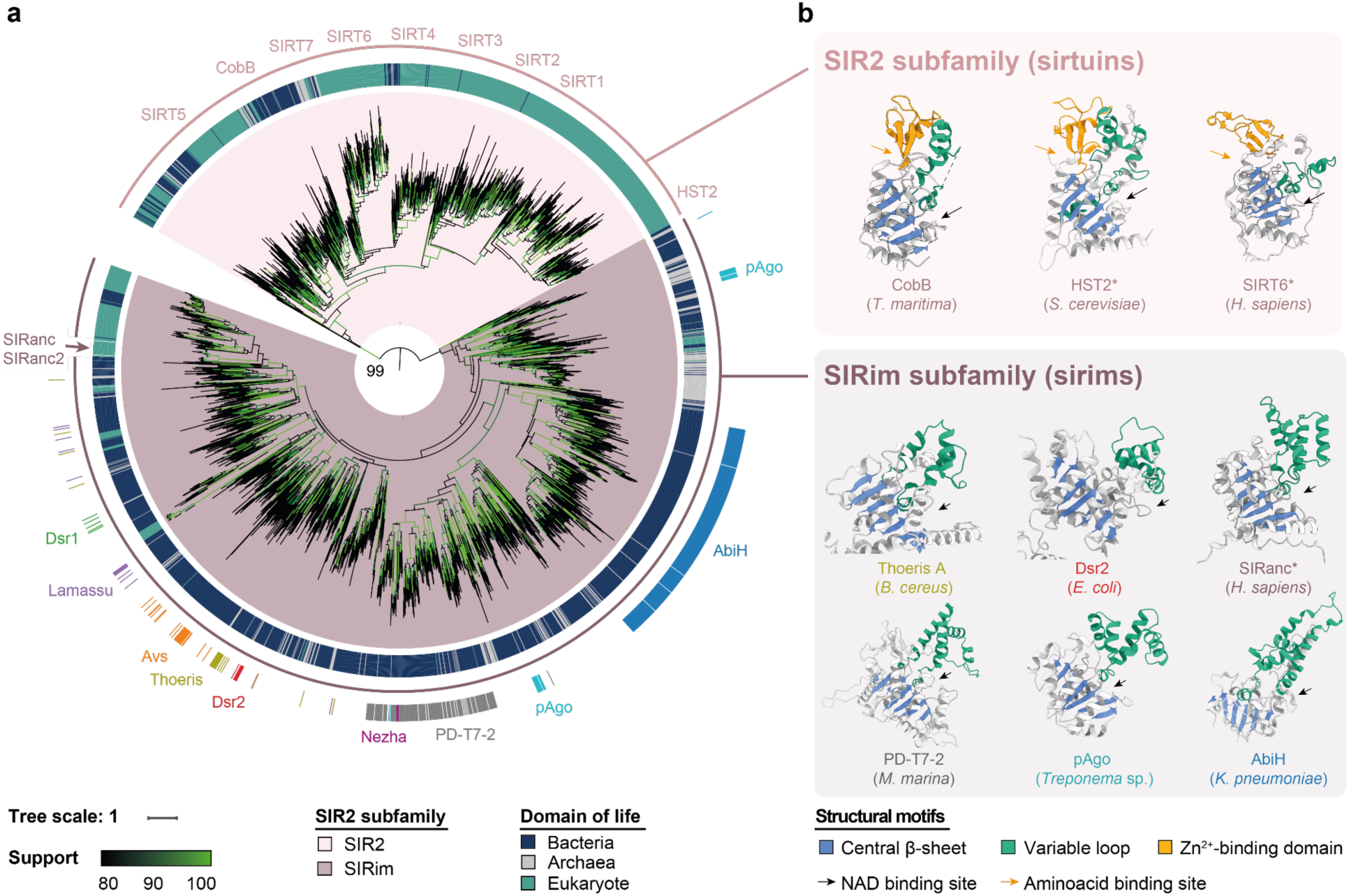
Human proteins belonging to the monophyletic group of SIRim proteins are associated with bacterial antiphage defence proteins. **a.** Phylogenetic tree of SIR2 family proteins across the tree of life. A total of 2,723 sequences representative of all the SIR2 domain-containing proteins identified through homology search were used to build the tree (midpoint rooted). SIR2 proteins cluster into two subfamilies based on amino acid sequences, namely SIR2 (inner ring, beige) and SIRim (inner ring, brown). The domain of life from which each protein originates is provided in the central ring (blue: bacteria, grey: archaea and green: eukaryotes). For proteins identified in bacterial organisms, we assessed if the protein is present in a known antiphage defence system and indicate its name (outer ring) (*21*). The location of human sirtuins (SIRT1-7), of SIRanc and SIRanc2, as well as of other proteins exemplified in Fig 1b are indicated. Tree branch support values are assessed using UltraFast Bootstrap and are indicated in shades of green. Support is also specified on the branch delimiting the two subfamilies. b. Structures of selected SIR2 proteins from the SIR2 (top, beige) and the SIRim subfamilies (bottom, brown). Three sirtuins (CobB (PDB: 2H2F), HST2 (PDB: 1SZC), SIRT6 (AF2: Q8N6T7)) and six sirims (Thoeris A (AF2: J8G6Z1, residues 1-279), Dsr2 (AF2: R8X3J8, residues 1-281), SIRanc (AF2: Q9BPY3), PD-T7-2 (AF2: I3Z638), pAgo (AF2: AIW90156), AbiH (AF2: A0A1V0M776)) are provided (PDB: Protein DataBank, AF2: AlphaFold2). For bacterial sirims, the name of the protein is coloured according to the antiphage system in which the protein is found. In ThsA, Dsr2, and pAgo, only the SIRim domain is shown while it is fused to additional domains in the original proteins. The central *β*-sheet of the Rossmann fold (blue) and the variable loop (green) are highlighted to underline the structural similarities between proteins. For each protein, the species of origin is indicated. A * after a protein name indicates that it is found in a eukaryotic organism.

To better characterise the two subfamilies, we mapped the proteins involved in bacterial immunity on the phylogeny. We first identified bacterial SIR2 proteins involved in documented antiphage systems with the program DefenseFinder (Fig 1a, outer circle) (*21*). All bacterial antiphage SIR2 proteins are located in one out of the two clades, suggesting that the entire clade may be involved in immunity (Fig 1a, brown clade). In contrast, all the bacterial and eukaryotic non-immune sirtuins are found in the other clade (beige clade) (*22*). Since DefenseFinder only identifies proteins whose antiphage action has been experimentally demonstrated, these predictions likely represent only a fraction of all existing antiphage systems. We thus sought to identify additional putative SIR2-containing antiphage proteins. To do this, we leveraged the tendency of antiphage genes to cluster within bacterial genomes, and analysed the frequency with which known antiphage systems are found in the vicinity (+/- 20 genes) of SIR2 proteins (*18*, *23*, *24*). The genomic neighbourhood of bacterial SIR2 proteins from the first, brown clade shows a 12-fold enrichment in antiphage systems compared to the neighbourhood of SIR2 proteins from the second, beige clade (Fisher exact test, F=12.40, p<10^-3^, Fig 1a & S1). Such bipartite distribution strongly suggests that the majority of the bacterial proteins from the first clade, but not from the second clade, have an immune function (*18*, *23*, *24*). This phylogenetic analysis led us to divide the SIR2 family of proteins into two subfamilies with seemingly distinct functions: i) sirtuins, which have a classical SIR2 domain and are involved in non-immune functions (Fig 1a, beige clade) and ii) sirims, equipped with a SIRim domain (for **SIR** and **im**munity, described hereafter), present in validated and predicted antiphage proteins, as well as in some eukaryotic proteins (Fig 1a, brown).

Analysing the experimental and predicted structures of SIR2 proteins sampled across the phylogeny reveals an overall conservation of the SIR2 domain topology, including the presence of a Rossmann fold with a 3-layer *α*/*β*/*α* sandwich. The central *β*-sheet contains a core of six *β*-strands, with accessory *β*-strands in a subset of clades (Fig 1b, blue). A variable *α*-helical loop is located downstream of the first *β*-strand (Fig 1b, green) (*25*). Sirtuins and sirims from bacteria, archaea and eukaryotes show conservation of the NAD^+^ binding pocket, present at the junction between the Rossmann fold and the variable loop, as well as of catalytic residues involved in NAD^+^ degradation (Fig S2a) (*16*, *17*, *20*, *26*). Structural differences however exist between sirtuins and sirims. The Rossmann fold of sirtuins, bearing a classical SIR2 domain, contains a *β*-sheet with exactly six *β*-strands, while seven or more strands are usually observed in sirim proteins. Sirtuins utilise a Zn^2+^ coordination site that allows binding of substrate amino acids of target proteins. Such a site is systematically absent from sirims, as is the tetrad of cysteines responsible for interactions with Zn^2+^ (Fig 1b, orange & Fig S2b). This suggests that proteins of the sirim clade are unable to bind acetylated amino acids, contrary to classical sirtuins (*16*, *26*). Overall, this structural and sequence analysis is congruent with the phylogenetic partition and suggests that the subfamilies of SIR2 proteins could have different functions.

We assessed the diversity of SIR2 proteins in humans by performing homology detection and identified 9 proteins. Seven of these are sirtuins, well-documented to regulate histone deacetylation (*15*, *22*, *27*). The two others, FAM118A and FAM118B, belong to the SIRim subfamily and have poorly-documented functions, with no known immune role (*28*). The evolutionary relationship between the FAM118 family of proteins and antiphage systems of bacteria led us to hypothesise that they may be involved in immunity. FAM118B was renamed SIRanc (sɪˈrɑːns) and FAM118A SIRanc2, after their **anc**estral **SIR** domain.

### SIRanc is involved in the mammalian TLR pathway

To gain insights into the function of SIRanc protein, we analysed the genomic localisation of *SIRanc* gene. In bacteria, genes coding for proteins of a given pathway tend to colocalise within genomes, displaying functional synteny. Although less common, multiple occurrences of a functional syntenic organisation have been documented in eukaryotes (*7–9*, *29*). In 73% of metazoans, including humans, *SIRanc* is located two genes away from the gene encoding for TIRAP (Fig 2a). TIRAP carries a TIR domain and contributes to TLR2 and TLR4 signalling. Upon ligand-induced TLR activation, TIRAP’s homotypic interaction with the cytosolic TIR domain of the receptor allows for immune signal transduction and production of proinflammatory cytokines (*30*, *31*). Synteny between the *SIRanc* and *TIRAP* genes suggests that SIRanc may be involved in the TLR pathway.

**Figure 2.**
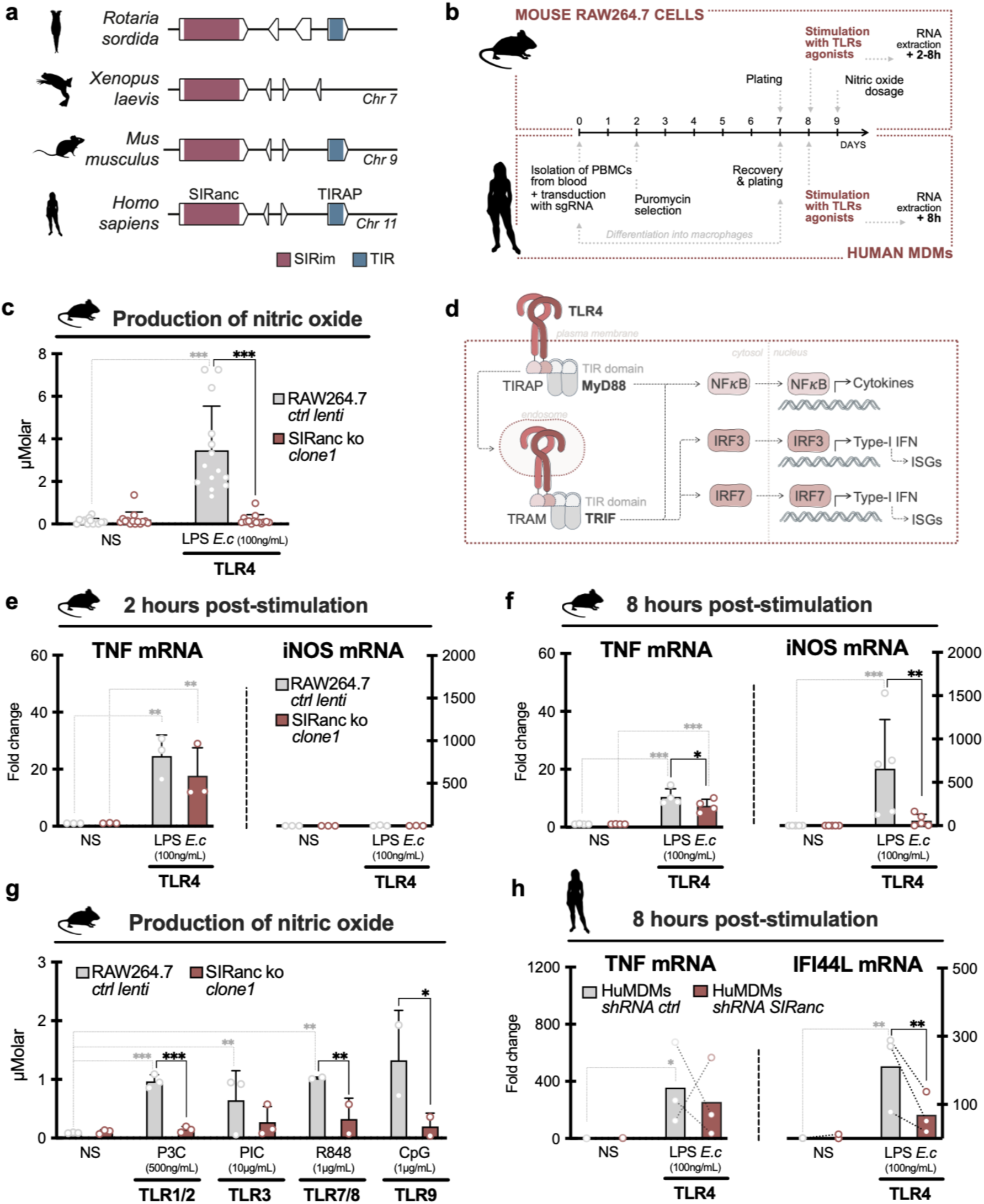
SIRanc contributes to TLR signalling. **a.** Examples of *SIRanc* genomic loci in animal genomes (metazoa). *SIRanc* (pink) colocalizes with *TIRAP* (blue) in 73% of metazoa. In *Xenopus laevis*, the African clawed frog, *TIRAP and SIRanc* are located on the same chromosome, but encoded at different loci. **b.** Experimental setup using murine RAW264.7 macrophage-like cells and human monocyte-derived macrophages (huMDMs) to probe the activation of the TLR pathway. **c.** Wild type or SIRanc KO murine macrophages were stimulated with LPS from *E. coli* to activate TLR4, and nitric oxide production was measured at 24h. **d.** Schematic of TLR4 signalling in macrophages. **e-f.** Wild type or SIRanc KO murine macrophages were stimulated with LPS and levels of TNF and iNOS transcripts were measured at 2h and 8h by RT-qPCR. **g.** Wild type or SIRanc KO murine macrophages were stimulated with Pam3CysSK4, Poly(I:C), R848 or CpG, agonists of TLR1/2, TLR3, TLR7/8 and TLR9, respectively, and nitric oxide production was measured at 24h. **h.** Human monocyte-derived macrophages stably expressing a control shRNA, or a shRNA downregulating the levels of SIRanc transcripts, were stimulated with LPS. Levels of TNF and IFI44L transcripts were measured at 8h by RT-qPCR. **c, e, f, g, h.** Each dot represents an independent experiment (**c, e, f, g**) or an independent human donor (**h**) (mean of technical replicates). Bars correspond to the mean and error bars to the standard deviation of all the independent experiments/donors. Statistical analyses were performed using two-ways ANOVA and p-values were classified as * p<0.05; ** p<0.01; *** p<0.001. Grey stars correspond to comparisons within one group and dark stars correspond to comparisons between groups.

Analysis of publicly available datasets indicate that *SIRanc* is broadly expressed in tissues and is enriched in macrophages (Fig S3a). The latter are essential immune cells that detect pathogens through pattern recognition receptors, including TLRs. TLR activation leads to an inflammatory and antimicrobial responses, as well as the priming of adaptive immunity. To address the role of SIRanc in the TLR pathway, we implemented a loss-of-function approach by knock-out in RAW264.7 murine macrophage-like cells (*ΔSIRanc* cells, Fig S3b-d). We initially assessed the TLR response by stimulating cells with purified LPS from *E. coli*, an agonist of TLR4, and measuring nitric oxide (NO) production (an antimicrobial compound). In wild type macrophages, LPS stimulation leads to a dose-dependent production of NO (Fig 2b-c & S3e). In *ΔSIRanc* macrophages, NO production is abrogated regardless of the LPS dose (Fig 2c & S3e). We controlled *ΔSIRanc* macrophages viability and observed no defect (Fig S3f). To verify that NO production is not mediated by receptor expression deficiency, we also verified that *ΔSIRanc* cells express wild type levels of TLR4 (Fig S3g). We additionally excluded LPS degradation in the absence of SIRanc by showing that both wild type and *ΔSIRanc* cells trigger pyroptosis in response to cytosolic LPS sensing through inflammatory caspases (Fig S3h) (*32*). The abrogation of NO production upon LPS stimulation in *ΔSIRanc* macrophages indicates that this protein plays a role in the TLR4 pathway.

We then aimed to determine whether SIRanc participates in a specific branch of the TLR4 pathway. Activation of TLR4 by LPS triggers signalling via two successive, distinct pathways which rely on specific adaptors, MyD88 and TRIF (Fig 2d). First, MyD88 activation triggers the upregulation of cytokines and interleukins. Among these, tumor necrosis factor (TNF) can be detected at 2h post-stimulation. Later, the TRIF-dependent pathway triggers the upregulation of interferon-stimulated genes (ISGs) as well as of the enzyme which synthesises NO (iNOS), observable at 8h post-stimulation (*33*, *34*). To evaluate the involvement of SIRanc in both branches, we monitored transcriptional changes of TNF, iNOS and ISGs (Rsad2) at 2h and 8h after TLR4 activation. As expected, in wild type macrophages, stimulation with LPS leads to the transcriptional upregulation of TNF at 2h and the upregulation of iNOS and Rsad2 at 8h (Fig 2e-f & S4a-b). At 2h, *ΔSIRanc* macrophages produce levels of TNF comparable to WT cells (Fig 2e & S4a). In contrast, at 8h, *ΔSIRanc* macrophages display lower levels of Rsad2 and iNOS compared to WT (Fig 2f & S4b). This is consistent with the lack of measurable NO at 24h observed in Fig 2c. These results demonstrate that SIRanc is involved in the late TLR4 signalling pathway in murine macrophages.

In addition to TLR4, it is well documented that other TLRs trigger upregulation of ISGs and production of NO (*35–37*). To interrogate if SIRanc is also involved in signalling downstream of these receptors, we stimulated wild type and *ΔSIRanc* macrophages with agonists of TLR1/2, TLR3, TLR7/8 and TLR9. In every case, lack of SIRanc translates into impaired NO production at 24h, highlighting the general role of SIRanc downstream of TLR stimulation (Fig 2g & S4c).

To assess the role of SIRanc in human innate immunity, we generated primary human monocyte-derived macrophages (huMDMs) knocked-down (KD) for SIRanc expression (Fig S5a). Consistent with our observations in murine macrophages, downregulation of SIRanc expression does not alter the production of TNF upon LPS stimulation, but significantly impairs the transcriptional upregulation of the ISG IFI44L (Fig 2h). We observed similar trends on other transcriptional readouts, as targets expected to be mostly MyD88-dependent (IL-8) are not affected by SIRanc downregulation (Fig S5b). On the other hand, the upregulation of TRIF-dependent ISGs (RSAD2 and MX1) is impaired in huMDMs SIRanc KD (Fig S5c). Concordant with the results obtained in murine macrophages, this data indicates that the activity of SIRanc is required for the late TRIF-dependent — but not early — TLR4 response in huMDMs.

Contrary to other TLRs, activation of TLR5 by the bacterial product flagellin only results in MyD88-dependent production of inflammatory cytokines. We assessed the transcriptional levels of IL-8, one of these inflammatory cytokines, in response to flagellin. We observed no difference between SIRanc KD and control huMDMs, consistent with the results obtained with LPS stimulation (Fig S5a-c). Altogether, these data document that SIRanc plays a pivotal role in ISG and NO production downstream of TLR stimulation in mice and human cells, demonstrating its role in mammalian innate immunity.

### SIRanc is an NADase

Sirims from bacteria and animals are evolutionarily related proteins that participate in immunity. Bacterial sirims perform an antiphage function by processing cellular NAD^+^ (*12*, *20*). To investigate the mechanism behind SIRanc inflammatory properties, we assessed whether SIRanc possessed NAD^+^ processing capabilities. Recombinant human SIRanc was purified and mixed *in vitro* with ε-NAD, an NAD^+^ analog whose degradation can be monitored using a fluorescence-based assay in which the fluorescent signal is proportionate to the amount of degraded ε-NAD. In the presence of ε-NAD alone, human recombinant SIRanc did not demonstrate detectable ε-NAD degradation activity (Fig 3a). This does not exclude the possibility that SIRanc requires an activation trigger, such as a protein-protein interaction, to initiate NADase activity. In bacteria, activation of a homologous sirim, ThsA, depends on oligomerization and formation of stable, enzymatically active protein filaments (*20*). Similarly, we hypothesised that SIRanc may be activated through oligomerization. To assess this, we introduced the molecular crowding agent polyethylene glycol (PEG) as a means of artificially stimulating protein-protein interactions *in vitr*o. In the presence of PEG, SIRanc demonstrated degradation of ε-NAD (Fig 3a).

**Figure 3.**
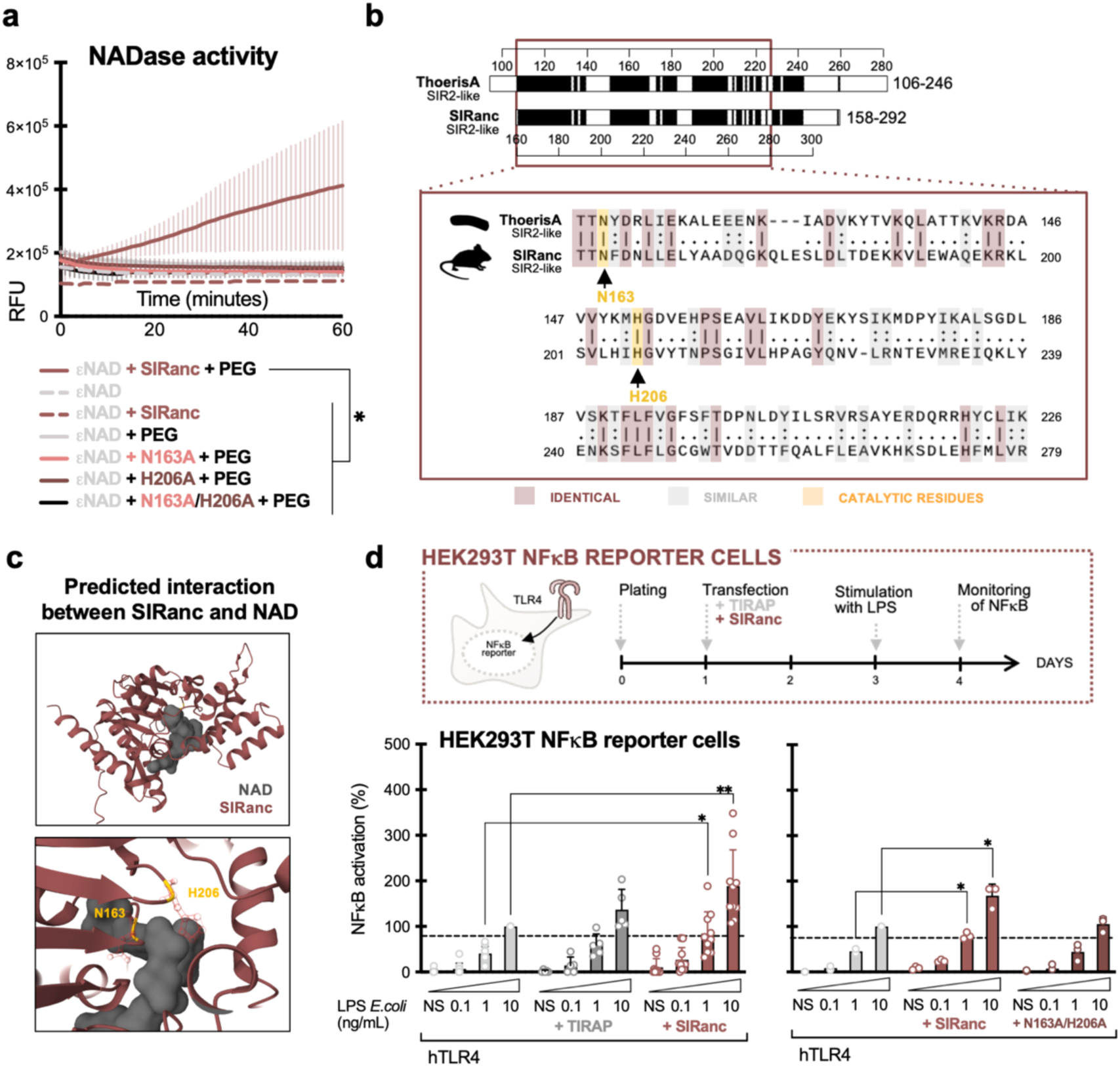
SIRanc is an NADase. **a.** Enzymatic NADase activity assay of purified recombinant SIRanc (wild type, N163A mutant, H206A mutant, N163/H206A double-mutant), incubated with ε-NAD, with or without PEG 400 over time. **b.** Local alignment of the amino acid sequences of the SIRim domains of human SIRanc and bacterial ThsA. Identical and similar residues are highlighted (red and grey, respectively), as well as the catalytic residues of ThsA (from (*12*), yellow). **c.** AlphaFold prediction of SIRanc (red) interacting with NAD^+^ (grey), with predicted interacting residues N163 and H206 (yellow). **d.** HEK293T-hTLR4 NFkB reporter cells were transiently transfected with plasmids coding for TIRAP, SIRanc wild type or SIRanc N163A/H206A double-mutant and were stimulated with LPS from *E. coli.* NFkB activation was measured after overnight stimulation. Each dot represents an independent experiment (mean of technical replicates). Bars correspond to the mean and error bars to the standard deviation of all the independent experiments. Statistical analyses were performed using two-ways ANOVA and p-values were classified as * p<0.05; ** p<0.01; *** p<0.001.

To clarify the similarity of SIRanc’s catalytic mechanism to that of other sirims, we investigated the involvement of homologous catalytic residues. The NADase activity of bacterial ThsA can be abolished by introducing alanine substitutions for the conserved catalytic residues N112 and H152 (*12*, *20*). To generate a catalytically inactive SIRanc, we mapped these conserved residues on the human sequence, confirming the residues’ putative homologous positions in an Alphafold2 model, and constructed N163A and H206A single and double mutants (Fig 3b-c). All predicted catalytically inactive SIRanc mutants had no detectable NADase activity, irrespective of the presence of PEG (Fig 3a). In summary, we document that *in vitro*, SIRanc is only active in the presence of a molecular crowding agent suspected to induce oligomerization, which is putatively responsible for activation of enzymatic NADase activity.

We next evaluated if SIRanc’s role in the TLR pathway depends on its NADase activity. Human embryonic kidney cells (HEK293T) engineered to express TLR4 and co-receptors, as well as a colorimetric reporter for NFκB-driven inflammation, can be utilised to monitor TLR4 signalling after LPS stimulation (Fig 3d). Ectopic expression of proteins can be achieved in this system by plasmid transfection, as exemplified by TIRAP, which induces a moderate, non-significant, increase in TLR4 activation (Fig 3d). Expression of wild type SIRanc induces an average two-fold increase in LPS-driven TLR4 signalling and NFkB activation, which is absent when the catalytically inactive version of SIRanc is expressed instead (Fig 3d). These results demonstrate that SIRanc participates in TLR signalling through its NADase activity.

### SIRim proteins are diverse and widespread among eukaryotes

We then investigated the evolutionary history of the SIRanc lineage. To do so, we queried Eukprot, a database containing genomes from 985 species spanning all major documented eukaryotic lineages, using the SIRim protein profile constructed in Fig 1 (*38*). SIRanc homologs are detected in a total of 35 diverse species including the freshwater protist *P. lacustris,* the zebrafish (*D. rerio*), the slime mold (*D. discoideum*), the Australian ghostshark (C. *milii*), and the mallard (*A. platyrhynchos*, Table S4). SIRanc homologs are distributed in a wide range of eukaryotic taxa suggesting that the acquisition in eukaryotic organisms is very ancient (Fig 4a-b & S6, Clade A). More precisely, the presence of SIRanc homologs in many genomes of opisthokonts, a eukaryotic taxon that includes fungi, choanoflagellates and animals (metazoa) suggests that the acquisition event occurred prior to their last common ancestor, more than 1.4 billion years ago, and that major classes of opisthokonta such as insects have lost the gene (Fig 4a-b, S6-S9; Table S4). We also observed that most amniotes, including mammals, possess two paralogs of SIRanc (namely SIRanc and SIRanc2), contrary to the rest of vertebrates (e.g. batrachia or bony fishes) which encode for only one SIRanc homolog (Fig S9). This indicates that SIRanc has undergone a duplication event, resulting in two copies of the gene. This occurred prior to the last common ancestors of amniota, approximately 320 million years ago (Fig S9).

**Figure 4.**
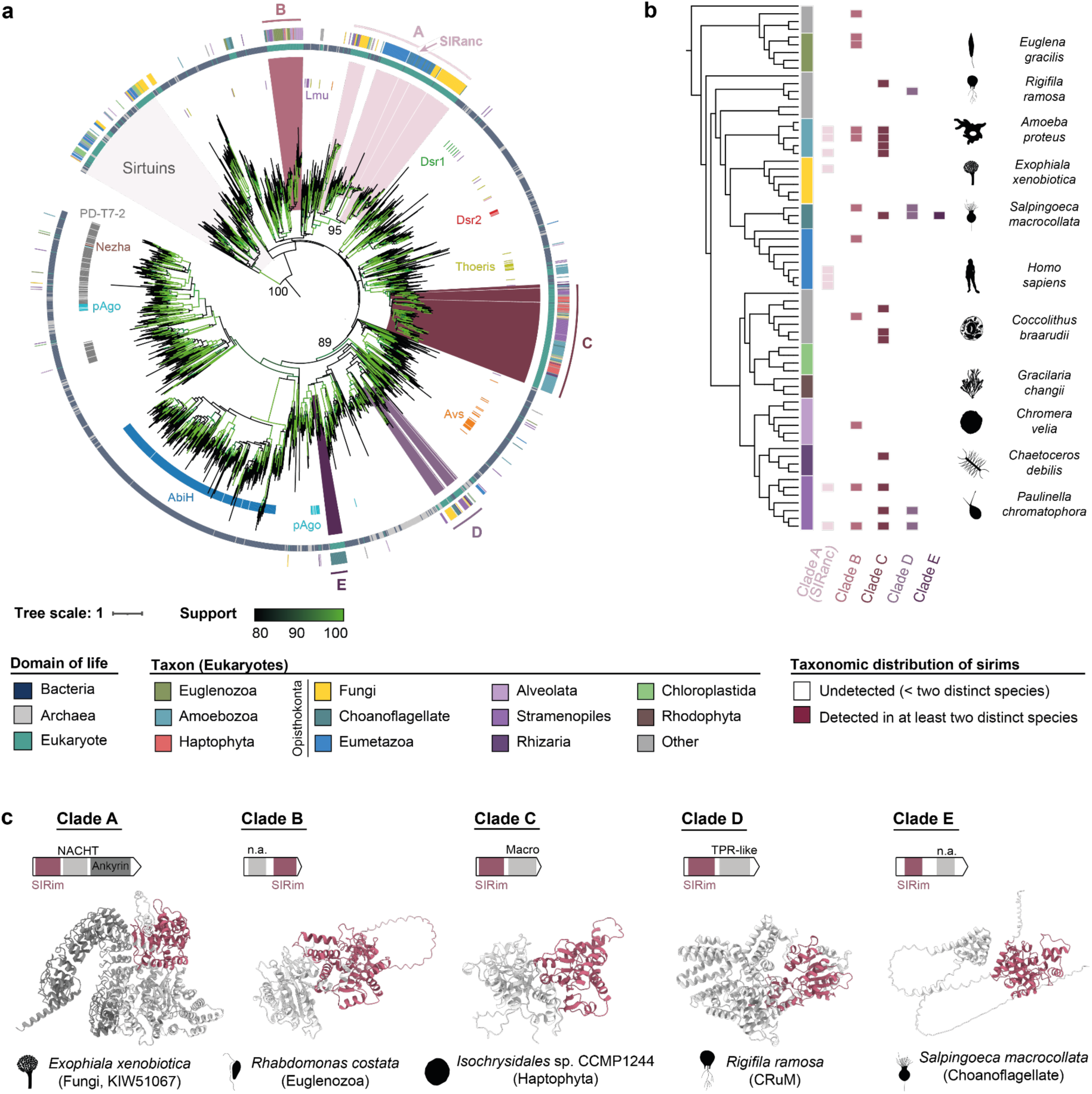
SIRanc is part of a large family of sirims in Eukaryotes. **a.** Phylogenetic tree of all identified SIRim domain-containing proteins across the tree of life. Representative sequences of all detected bacterial, archaeal, and eukaryotic sirims (*n*=2,266) are aligned on their SIRim domain and a phylogenetic tree was built from the multiple sequence alignment. The domain of life of each protein is indicated in the central ring (blue: bacteria, grey: archaea, green: eukaryotes) and the specific taxon is indicated for eukaryotic sequences in the outer ring. Antiphage systems are annotated using DefenseFinder for bacterial sirims (inner ring) (*21*). The five major eukaryotic clades of sirims (clades A to E) are coloured in shades of purple (inner ring) and within each clade, specific sequences are exemplified in Fig 4c. Branch support is assessed using UltraFast Bootstraps and is indicated in shades of green. Support values are indicated on key branches. **b.** Taxonomic distribution of SIRim proteins in Eukaryotes. The phylogenetic tree of The Comparative Set (TCS) from Eukprot includes 196 species from all major known eukaryotic phyla (*38*). The taxonomic distribution of each of the five eukaryotic SIRim protein clades is displayed. SIRim proteins detected in species present in the Eukprot v3 database (985 species) but not in the TCS (196 species) are assigned to their closest relative present in the TCS tree. Sirim presence in a specific clade is reported only when at least two distinct species of the clade are identified as encoding a sirim gene in their genome. The tree is midpoint rooted. The tree is also pruned, and branch lengths are ignored for visualisation purposes (see Fig S6 for a complete version of this tree). For each major eukaryotic taxon, a logo of one example species encoding a *sirim* gene is provided. **c.** Examples of sirims found in eukaryotic organisms. Sirims are sampled from each of the five major eukaryotic sirim clades and are exemplified (Fig 4a & Fig S7-S8). For each sequence, the domain architecture as well as the AlphaFold2 model is provided. The SIRim domain is coloured in pink and additional, fused, domains are indicated in grey. Whenever the specified domain could be characterised using HHpred, its name is provided. NACHT (PF05729), TPR-like (PF14938), n.a.: domain not identified.

Beyond the lineage of SIRanc homologs (Clade A), our phylogenetic analysis uncovers four additional lineages of eukaryotic sirims (Clades B to E) in 189 different species. In total, 19.2% of the eukaryotic genomes we queried encode a SIRim protein (Fig 4a & S6-S8, Table S4). Clades A to C are distributed among a wide diversity of eukaryotes, with for example clade C, regrouping 83 species. Conversely, clade D and E are limited to a small number of eukaryotes, with clade E detected only in choanoflagellates (Fig 4b & S6, Table S4). The interspersed distribution of the five eukaryotic lineages within prokaryotic clades in the phylogenetic tree strongly suggests that the five clades of eukaryotic sirims arose independently during the history of eukaryotes. Interestingly, these lineages branch with prokaryotic sirim proteins belonging to different antiphage systems (clades B and C with ThsA-like, and clades D and E in pAgo-associated sirims). These multiple acquisition events were followed by frequent gene losses, explaining the patchy distribution of sirims across eukaryotes (Fig S7 & S8).

Finally, we sought to gain insights regarding the biological function of eukaryotic sirims from the five identified clades. Strikingly, we observed that the SIRim domain is frequently fused to additional domains which are often observed in immune proteins. Clade A, for example, contains fungal and animal sirims with an N-terminal SIRim domain, a central nucleotide-binding domain (such as NACHT or NB-ARC), and C-terminal repeats (*e.g.,* ankyrin or tetratricopeptide). This tripartite architecture is typical of nucleotide-binding domain, leucine-rich repeat-containing (NLR) proteins, which are immune receptors key to detecting a variety of infection cues in mammals or plants as well as to inducing programmed cell death upon mating of incompatible germlings in fungi (Fig 4c, Clade A) (*39–42*). In clade C, sirims are typically composed of a SIRim domain fused to a Macro domain (Fig 4c). This mirrors the domain architecture of ThsA from the type II Thoeris antiphage system (*43*, *44*). The genetic architectures of sirim proteins thus suggest that they play an immune role in a wide diversity of eukaryotes.

## Discussion

In this work, we demonstrate that the widespread SIRim domain involved in antiphage defence also participates in eukaryotic immunity. We uncover the role of SIRanc, a mammalian innate immune protein with a SIRim domain involved in late signalling downstream of TLRs, responsible for ISG upregulation and iNOS production in macrophages. Such proinflammatory role is dependent upon SIRanc’ NADase activity.

Given the vast body of literature on TLR pathways, one may be surprised that such a pivotal protein was not discovered sooner, through the implementation of a CRISPR screen or otherwise. One such CRISPR-Cas screen was performed to identify members of the TLR cascade, but measured TNF levels as a read-out of TLR4 activation (*45*). TNF is produced mainly through early TLR4 signalling, and our data indicate that its production is independent of SIRanc, explaining why it was not identified in that study. A recent publication also mapped members of the myddosome, a MyD88-dependent supramolecular structure responsible for signalling downstream of TLR activation (*46*). SIRanc was not identified in this work. In line with SIRanc’s link to TRIF, it suggests that its mechanism of activation may not rely solely on interactions with myddosome components.

While our results document the importance of SIRanc within the TLR transduction cascade, follow-up studies will focus on delineating the regulatory mechanisms governing its enzymatic activity. One key question relates to the downstream effectors acting after SIRanc. We hypothesise that SIRanc may produce an inflammatory mediator from NAD^+^ degradation. Data from the literature involve transient receptor potential melastatin-related (TRPM) in the inflammatory response driven by TLR4. For example, TRPM2 is activated by the binding of products of NAD^+^ degradation, including ADPR and cADPR, to its SLOG domain. This leads to the production of proinflammatory cytokines such as IL-1α and CXCL2, as well as the upregulation of iNOS, but does not affect TNF secretion (*47*, *48*). Whether TRPM2, other TRPM channels, or other proteins can detect SIRanc products of NAD^+^ degradation remains to be explored.

The SIRim domain is involved in multiple antiphage systems, protecting bacteria against viral infection. In mammals, it is integrated within the TLR pathway, which detects viruses, but also bacterial infections. There is thus a diversification of this domain’s protective role in eukaryotes. Conserved antiphage domains appear tailored to clade-specific immunity, in link with selective pressures arising from eukaryote and/or clade-specific pathogens. Such adaptability was observed for additional conserved immune proteins, such as cGAS/cGLRS and viperins (*8*, *9*, *49*). At the molecular level, the adaptation of the SIRim domain to the context of eukaryotic innate immunity translates into a shift in its biochemical activity: from a strong NADase in bacteria that depletes NAD^+^ and triggers cell death, to a weaker NADase activity in humans, that does not trigger cell demise but probably participates in signal transduction.

Beyond humans and animals, we uncovered hundreds of sirims in a wide diversity of eukaryotes, including fungi, amoeba, alveolata, and diatoms. Other domains of antiphage origin also show conservation across a wide range of eukaryotes. TIR domains participate in immune signal transduction within the Toll receptor of insects and as part of mammalian TLRs (*11*). Homologs of cGAS, coined cGAS-like receptors (cGLRs), are conserved across metazoans and participate in immunity in species such as coral and oyster (*49–51*). Based on SIRim-containing proteins being frequently fused with domains involved in immunity, we predict that sirims could participate in host defence in many of these – often non model – organisms. We have, for example, identified SIRim homologs in *Chaetoceros* diatoms, which are keystone organisms in marine ecosystems. Studying immunity in these organisms could improve our understanding of how host-pathogen interactions shape ecosystems.

To account for the ever-growing number of immune members conserved across domains of life, we recently proposed the concept of ancestral immunity (*52*). Ancestral immune modules are protein or protein domains evolutionary conserved between prokaryotes and eukaryotes, involved in immunity in both kingdoms. We show in this work that, akin to PNP, NACHT or TIR, SIRim corresponds to an ancestral immune module. Conservation across kingdoms can be harnessed for mechanistic studies, by interrogating the module’s molecular features in multiple species in parallel (e.g., bacteria and humans). Whether SIRanc plays a role in human pathology, and can be harnessed for therapy, will be the topic of future investigation.

## Acknowledgments

We are grateful to members of the MDM lab, U1284, U932 as well as mentors and friends for their useful comments on earlier versions of the manuscript. We thank C.W. for the kind gift of cell lines and S.H.L., A.W., and E.H. for cloning assistance. Some of the bioinformatic analyses were performed on the Core Cluster of the Institut Français de Bioinformatique (IFB) (ANR-11-INBS-0013). We also thank the team managing the Phylopic database of logos as well as all the contributors (https://www.phylopic.org/); the freely shared content helped us design certain figures of the paper.

## Author contributions

A.B and E.Z.P conceptualised the project. All the authors designed experiments and analysed data. D.B., H.V., E.M.Q.E., V.R., J.C. and V.H.T. performed data curation and formal analysis. D.B., E.M.Q.E., B.R.M., V.R. and V.H.T. performed experiments. H.V. ran all the bioinformatic analysis. All authors contributed to writing of the manuscript. A.B., E.Z.P., P.B. and B.R.M. acquired funding.

## Data and materials availability

All the data, including phylogenetic trees, HMM profiles and protein structures generated with AlphaFold, are available at https://gitlab.pasteur.fr/mdm-lab/siranc.

## Materials and Methods

### Database of eukaryotic, archaeal and bacterial genomes

We downloaded 4,616 Eukaryotic genomes from 2,407 species and 22,803 Bacterial complete genomes representing 7,231 species from the Genbank database (https://ftp.ncbi.nlm.nih.gov/genomes/all/GCA/) in January 2022 (*53*). 2,339 Archaeal genomes were also downloaded from the Genome Taxonomy DataBase (GTDB) in January 2023 and concatenated to them 496 inhouse Asgard Archaea genomes resulting in a database of 2,835 Archaeal genomes (*54*). This resulted in a database of 30,254 genomes spanning across the three domains of life.

### Systematic detection of SIR2 and SIR2-like proteins using iterative profile HMM search

An initial, loose, detection of SIR2 family proteins was performed by building a general profile HMM modelling the SIR2 family of proteins. This was done by aligning known SIR2 homologs including (i) known eukaryotic homologs of sirtuins (e.g. orthologs of *H. sapiens* SIRT1 to SIRT7) as well as (ii) homologs of bacterial antiphage proteins containing a SIR2 domain and identified by running DefenseFinder (v1.2.3, default parameters) on our database of genomes (*21*). SIR2 genes from the following systems were included in the selection of antiphage SIR2 proteins: Thoeris, Dsr1, Dsr2, PD-T7-2, Nezha, Avs, AbiH, Lamassu and prokaryotic Argonautes. This yielded a set of *n=4,302* sequences which were aligned using Muscle v5.1 (option -super5) and a profile HMM was built using the hmmbuild command from HMMER (version 3.3.2) (*55*, *56*). This initial SIR2 family profile was queried against our database of genomes using hmmsearch (HMMER package) which yielded *n=57,911* hits satisfying the inclusion criteria (e-value < 1e-5, bit score > 20, query coverage > 60% and 100 < hit length < 2500 amino acids) in bacteria, archaea and eukaryotes (Table S1). These proteins were clustered using MMseqs2 v13.45111 easy-cluster (30% identity, 80% coverage), representative sequences were aligned using Muscle v5.1 and the alignment was trimmed using Clipkit v1.3.0 (options -m gappy -g 0.99) (*57*). An initial maximum likelihood phylogenetic tree was built from the trimmed alignment using IQTree v2.2.0 using the LG+G4 substitution model (*58*). Maximum likelihood optimization was performed for 1000 iterations (data not shown).

A refined detection of SIR2 proteins was then performed per SIR2 subfamily (SIR2 and SIRim separately) using an iterative profile HMM search against our database of bacterial, archaeal and eukaryotic genomes. This detection was performed to get a curated set of proteins per subfamily, to be used in all the remaining analyses. First, FAM118A and FAM118B orthologs were fetched from the NCBI Orthologs database (https://www.ncbi.nlm.nih.gov/gene/79607/ortholog), by filtering out proteins designated as “PREDICTED” or “LOW QUALITY” leading to a set of *n=453* FAM118 orthologs from Vertebrate organisms. Proteins were aligned using Muscle v5.1 (option -super5) and a profile HMM of FAM118A and FAM118B orthologs was constructed from the Multiple Sequence Alignment (MSA) using hmmbuild from the HMMER package (*55*, *56*). This initial profile HMM was queried against the database of Eukaryotic proteins only in order to identify FAM118 homologs in this domain of life. This search (“Round 1”) led to obtaining *n=751* hits among Metazoa and Fungi satisfying the including criteria (e-value < 1e-05, 200 < length < 600 aminoacids). Hits were aligned using Muscle v5.1 (option -super5) and and a new profile HMM was built to search the database of eukaryotic genomes for three additional round (“Round 2” to “Round 4”) using HMMER, until no additional SIR2-domain containing proteins was identified in eukaryotes (Table S3) (*55*, *56*). At each round, among all the profile HMM hits, only those satisfying the inclusion criteria were included in the list of valid hits and were included in the profile HMM for the subsequent profile HMM search (Table S3). FAM118 homology search was then expanded in Bacteria and Archaea using four additional rounds of profile HMM search (“Round 5” to “Round 8”) against our database bacterial, archaeal and eukaryotic genomes. At each round, only hits satisfying the inclusion criteria were included to the list of valid hits (Table S3). Iterative search was stopped once no additional hits satisfying the including criteria were retrieved from the database, yielding *n=8,697* of sirims across domains of life. To this set of proteins were added the list of abiH homologs (*n=1,208*) which were retrieved by querying the PF14253 Pfam profile against our database of genomes using hmmsearch (--cut_ga option) (*56*). This led to a total of 9,905 hits including SIRanc/SIRanc2 (FAM118A/B) homologs in eukaryotes as well as proteins involved in known bacterial antiphage defense systems.

All deacetylating sirtuins were also retrieved from the database of genomes using a similar procedure of iterative profile HMM search. Initially, orthologs of all human sirtuins (SIRT1 to 6) were fetched on the NCBI Orthologs database and hits with “LOW QUALITY” or “PREDICTED” headers were filtered out. The set 3,861 human sirtuin orthologs was clustered using MMseqs2 easy-cluster at 98% identity and 98% coverage, representative sequences were aligned using Muscle v5.1 and a profile HMM of human sirtuin orthologs was built using hmmbuild (*55*, *56*, *59*). The obtained profile was iteratively queried against our database of eukaryotic genomes for three rounds (“Round 1” to “Round 3”) until no novel eukaryotic sirtuins were detected (Table S2). After round 3, hits satisfying the inclusion criteria (e-value < 1e-5, bit score > 110, query coverage > 80% and 200 < protein length < 1500 amino acids, *n=23,192*) were clustered at 50% identity / 80% coverage using MMseqs2 easy-cluster. Representative sequences (*n=1,944*) were aligned using Muscle v5.1, a profile HMM was built using hmmbuild and the profile was queried against bacterial, archaeal and eukaryotic genomes. Iterative homology search was performed across the tree of life for two additional iterations until no additional sirtuin homologs were detected, leading to a set of *n=46,623* detected sirtuin homologs satisfying the inclusion criteria (e-value < 1e-5, bit score > 80, query coverage > 60% and 200 < protein length < 2500 amino acids, (Table S2).

### Annotation of defence systems in bacterial genomes

To assess which SIR2 family proteins are involved in known antiphage systems, we extracted the genomic neighbourhood (twenty genes upstream, twenty genes downstream) surrounding each sirtuin and sirim identified in the refined detection and ran DefenseFinder v 1.2.3 with default parameters to identify which defence systems contains a sirtuin or a sirim (Fig 1a) (*21*). We also identified whether the genomic neighbourhood of bacterial sirtuins and sirims is enriched in defense systems as antiphage defense systems to cluster together in bacterial genomes (*23*, *24*). We measured whether at least one defense system occurs (Fig S1a) and as well as how many defense systems (Fig S1b) can be found in the genomic neighbourhood of sirtuins and sirims (20 genes upstream and downstream). The difference in the frequency of neighbouring defense systems between sirims and sirtuins was assessed using a two-sided Fisher exact test (Fig S1a) and the difference in the number of neighbouring defense systems was assessed using a two-sided Mann-Whitney test (Fig S1b), as implemented in the scipy python package (v1.8.0) (*60*).

### Phylogenetic inference

We first built a phylogenetic tree for the whole SIR2 family (including both SIR2 and SIRim subfamily proteins) using the proteins identified in the refined detection (Fig 1a). All identified sirtuins (*n=46,623*) and sirims (*n=8,697*) were clustered differentially in order to increase the representation of eukaryotic and archaeal proteins compared to their bacterial counterparts (*59*). Precisely, sirtuins were at 50% identity-80% coverage, bacterial sirims were clustered at 50% identity-80% coverage whereas archaeal and eukaryotic sirims were clustered at 90% identity-90% coverage. A total of 2,723 representative sequences (*n=947* sirtuins and *n=1,776* sirims) were aligned on their SIR2 domain using Muscle v5.1 (option -super5 -permut none), the alignment was trimmed using Clipkit v1.3.0 (options -m gappy -g 0.99) and was used to build a maximum likelihood phylogenetic tree using IQTree (*55*, *57*, *61*). The Q.pfam+G4 amino acid substitution model was selected using ModelFinder and the Bayesian Information Criterion(*62*). Maximum likelihood tree search was performed until convergence and branch support was assessed using 1,000 UltraFast Bootstrap trees (*63*).

We then built phylogenetic trees of all sirims identified in our database of genomes (Fig 4a, Fig S7 & S8). Prior to building a phylogenetic tree, in order to increase the representation of undersampled eukaryotic clades (e.g. rhizaria or alveolata), representative sequences of all identified sirims in the refined detection (*n=8,687*) were used to build a profile HMM that was queried against the Eukprot v3 database using hmmsearch (*38*). Hit satisfying the inclusion criteria (e-value < 1e-5, bit score > 55, query coverage > 60% and 100 < protein length < 2500 amino acids, *n=482* taking diverse isoforms of the same protein into account) among 205 genomes (over a total of 985 in the Eukprot v3 database) were further filtered to remove putative bacterial contaminations. To do so, all *n=482* Eukprot v3 hits were used as queries with Diamond v1.2.6 (option --fast) against the NCBI nr database of proteins (*64*). Eukprot v3 proteins having more than sequence 70% identity with any bacterial protein in nr were filtered out. This led to a curated set of *n=443* sirims detected in *n=189* different genomes from the Eukprot v3 database (Table S4). These curated hits were included in the set of all sirims used to build the tree. All sirims were clustered differentially depending on their domain of life using MMseqs2 easy-cluster : bacterial sirims were clustered at 50% identity-80% coverage whereas eukaryotic and archaeal sirims were clustered at 90% identity-90% coverage in order to increase their abundance in the tree compared to their bacterial counterparts (*57*). The set of representative sequences was manually cured by removing proteins lacking key amino acids in the GxG motif and in the xxNxD motif and gap-inducing proteins at these loci. Sirims found in the *Homo sapiens* genome were explicitly added to the set of representative sequences. This step led to a total number of *n=2,101* cured representative sequences of sirims. We also included a set of sirtuins from across the three domains of life to the tree of sirims to serve as an outgroup. To do so, *n=165* sequences were sampled from the set of representative deacetylating sirtuin sequences (clustered at 50% identity/80% coverage using MMseqs2 easy-cluster) were sampled from all three of domains of life and added to set of proteins included in the tree of sirims (*57*). The obtained representative sirtuins and sirims (*n=2,266*) were then aligned on their SIR2/SIRim domain using Muscle v5.1 (option -super5), the MSA was trimmed using clipkit (option gappy, 99%) and a maximum likelihood phylogenetic tree was inferred using IQTree v3 (*58*). The Q.pfam+G4 substitution model was selected via ModelFinder using the Bayesian Information Criterion (BIC), and the tree was optimised until convergence and branch support was assessed using 1,000 UltraFast Bootstraps (Fig 4a) (*62*, *63*). To assess if the topology of the phylogenetic tree of sirims is stable to perturbations of the input MSA, additional SIR2/SIRim domain MSA were generated using (i) MAFFT v7.505 L-INS-i algorithm (option --localpair) and (ii) Muscle v5.1 by permuting the guide tree when building the alignment (option -super5 -perm all). This procedure yielded five additional MSA of sirims which were trimmed using clipkit (-m gappy -g 0.99) and were used to build five phylogenetic trees with IQTree (same procedure as before) (*55*, *58*, *61–63*, *65*) (Fig S7). Another phylogenetic tree containing only eukaryotic sirims was built (Fig S8). All detected eukaryotic sirims were clustered at 90% identity and 90% coverage using MMseqs2 leading to a set of *n=485* representative sequences which were aligned using Muscle v5.1 (option -super5) together with *n=165* sirtuins from across the tree of life serving as an outgroup to root the tree. The alignment was trimmed using Clipkit (-m gappy -g 0.995) and the tree was built using the Q.pfam+G4 substitution model and maximum likelihood search was performed until convergence (*55*, *58*, *59*). Tree branch support was assessed using 1,000 generated UltraFast Bootstrap.

In all the phylogenetic trees of sirims, we looked for clades of eukaryotic sirims (Fig 4a-4b & Fig S6-8). The criteria to consider that a eukaryotic clade is valid are the following (i) the clade of eukaryotic proteins is well supported, (ii) the clade contains sequences from at least two distinct eukaryotic species, (iii) the genomes in which sirims are detected were produced in at least two distinct studies and (iv) the clades are stables to perturbations in the phylogenetic analysis (*66*). In the case where a clade of bacterial proteins forms a well-supported clade with the eukaryotic proteins and is basal to it, we consider that a putative Horizontal Gene Transfer (HGT) event from bacteria to eukaryotes is likely to have occurred (*66*).

To compute the taxonomic distribution of each lineage of sirims in eukaryotes, we analysed the set of sirims detected in the Eukprot v3 database (genomes from *n=985* species). As no phylogenetic tree was available for the full Eukprot v3 species, we plotted the distribution on the phylogenetic tree of The Comparative Set (TCS) from Eukprot (genomes from *n=196* species) (*38*). For each species in Eukprot v3, either the species is also present in TCS, or it is not. In the latter case, the species is assigned to its closest relative in Eukprot TCS using TaxonKit (*67*). For each species in TCS, we consider that sirims are detected if (i) sirims are detected in at least two Eukprot v3 different species represented by this species or (ii) the clade represented by the TCS species contains only one Eukprot v3 species and sirims are detected within it (Fig 4b & Fig S6). Divergence time between eukaryotic taxa was taken from the TimeTree 5 (http://www.timetree.org/) (*68*).

### Domain annotation and folding of eukaryotic sirims

We sought to annotate the protein domains of all detected eukaryotic sirims (Clades A to E, Fig 4c). Domain annotation was performed using the HHpred server (default parameters) against Pfam-A v36 and the Protein DataBank (PDB) mmCIF70 (*69*). Hits satisfying inclusion criteria (probability > 0.90) were considered valid. Proteins were also folded using AlphaFold2/ColabFold as implemented in the localcolabfold package v1.5.2 (https://github.com/YoshitakaMo/localcolabfold, single prediction using 12 optimization recycles) (*70*, *71*).

### RAW264.7 cells culture and agonist stimulation

RAW264.7 macrophage-like cells (initially purchased from ATTC) were kindly provided by Dr. Catherine Werts (Institut Pasteur, Paris, France). KO cell lines were generated using transduction with Lenti-CRIPSR-v2 containing sgRNA targeting fam118b or non-targeting control guides (Fig S3b & Table S5). Transduction was followed by puromycin selection, single cell cloning, and knock-out phenotype was confirmed using both PCR showing genomic deletions, and western blot confirming the total absence of protein expression (Fig S3c-d).

RAW264.7 cells (control and FAM118B KO) were cultivated in complete RPMI medium (Gibco) (10% FCS + Penicillin-Streptomycin) and were plated at 0.5 x 10^6^ cells/mL in 200µL/96-well plate or 1mL/24-well plate (TPP) the day before stimulation. Stimulation with TLRs agonists was performed the day after plating. Cells were stimulated for 2h-24h with LPS of *Escherichia coli* strain O111:B4 (LPS-EB, Invivogen), synthetic triacylated lipopetide (Pam3CysSK4, Invivogen), dsRNA analog poly(I:C) (PIC-HMW, Invivogen), flagellin of *Salmonella typhimurium* (FLA-ST UP, Invivogen), resiquimod (R848, Invivogen) or short synthetic single-stranded DNA (CpG ODN 1018, Invivogen). After stimulation, cells were frozen immediately for subsequent RNA extraction and supernatants were recovered for dosage of nitric oxide.

### Nitric oxide dosage

Nitric oxide (NO) production by macrophages was measured in fresh macrophages supernatants by the Griess reaction (Griess reagent system, Promega), according to the supplier’s recommendations.

### Propidium iodide integration and LDH release assay

RAW264.7 cells (control and FAM118B KO) were plated at 0.25 x 10^6^ cells/mL in 200µL/96-well plate the day before transfection. Transfection with LPS of *Escherichia coli* strain O111:B4 (LPS-EB, Invivogen) was performed using a transfection reagent (FuGENE, Promega) in OptiMEM (Gibco). Cells were then incubated overnight and LDH release in the supernatant was measured (CyQUANT cytotoxicity assay, Thermofisher) according to the manufacturer’s instructions. In parallel, the integration of 1µg/mL propidium iodide was measured by fluorescence.

### Primary human monocyte-derived macrophages

Plasmapheresis residues from healthy adult donors were obtained from the Établissement Français du Sang (Paris, France) in accordance with the Institut National de la Santé et de la Recherche Médicale (France) (INSERM) ethical guidelines. According to French Public Health Law (art L 1121–1-1, art L 1121–1-2), written consent and Institutional Review Board approval are not required for human non interventional studies.

Human peripheral blood mononuclear cells (PBMCs) were isolated from the blood of healthy donors using Ficoll-Paque (GE Healthcare) and CD14+ monocytes were then positively selected using magnetic beads (Miltenyi). Differentiation of the cells into macrophages was induced in RPMI medium (Gibco) with 5% FCS + 5% human serum AB (Sigma), penicillin-streptomycin (Gibco) with 50 ng/mL human macrophage colony-stimulating factor (M-CSF, Miltenyi) for 7 days.

For cell transduction, 10 x 10^6^ monocytes were plated in 10-cm non-treated cell culture dishes and treated with 5 mL of freshly-collected lentiviral particles (see below), in the presence of 8 mg/mL of protamine sulphate (Sigma Aldrich). Selection of the transduced cells was performed with 2 µg/mL of puromycin 48 hours post-transduction. Seven days post differentiation and transduction, human monocyte-derived macrophages (MDMs) were collected, plated and stimulated with agonists of TLRs as described above.

### shRNA-lentiviral particles production

HEK293-LTV cells were maintained in DMEM supplemented with 10% FCS, 100 U/mL penicillin/streptomycin, and 1% (V/V) of non-essential amino acids (Gibco). Cells were routinely sub-cultured. For lentiviral production, 7 x 10^6^ cells were seeded in T75 cm^2^ flasks and transfected the next day. Transfection mixes consisted of 4.1 mg of lentiviral genome, 3 mg of the packaging plasmid PSPAX2 and 1.25 mg of VSV-G. DNA was complexed with 7.5 mM of polyethylenimine (PEI) for 20 min and added to cells. Sixteen hours later, culture media was renewed, and the lentiviral particles allowed to produce and accumulate for an additional 30 hours, after which they were collected, filtered at 0.45 mm, and immediately used to transduce monocytes. The lentiviral vectors employed were; pLKO.1-puro Non-Mammalian shRNA Control Plasmid DNA (SHC002, Sigma) and pLKO.1-Puro shRNA-FAM118B (TRCN0000130258, Sigma).

### RNA extraction and RT-qPCR analysis of cytokines and ISGs

RNA extraction was performed on frozen cells using NucleoSpin RNA (Machery-Nagel), according to the supplier’s instructions. Reverse transcription was performed with SuperScript RT (Invitrogen). RT-qPCR was performed with Taqman Universal Master Mix (Applied Biosystems), according to the manufacturer’s instructions (Table S6). Fold change was calculated using the 2^-ΔΔCt^ method, with normalisation by internal control (GAPDH for mouse cells and HPRT for human cells) followed by normalisation by the non-stimulated condition.

### Plasmids

Plasmids were ordered from GenScript: pcDNA3.1 backbone (neomycin resistance) with either human (Hs) FAM118B, with a N-term FLAG-tag, or human (Hs) TIRAP, with an N-term HA-tag. Alanine substitution of HsFAM118B (N163A & H206A) was performed by site directed mutagenesis (Quick change XL, Agilent), according to the manufacturer’s instructions. All Minipreps and Midipreps were performed using endotoxin-free kits (Macherey-Nagel).

### TLR4-NFkB reporter systems

HEK293T-Blue-Null2 and HEK293T-Blue-hTLR4 (initially purchased from Invivogen) were kindly provided by Dr. Catherine Werts (Institut Pasteur, Paris, France). NFkB reporter HEK293T-Blue-Null2 and HEK293T-Blue-hTLR4 cells (stably transfected with human TLR4, MD2 and CD14) were cultivated in complete DMEM medium (Gibco) (10% FCS + Penicillin-Streptomycin). FAM118B and TIRAP overexpression was achieved by transfection of both cell lines with either 10 ng/well pcDNA3.1-HsFAM118B-Flag (or catalytic mutant) or 0.1 ng/well pcDNA3.1-HsTIRAP-HA in 12-wells plates (TPP), using lipofectamine (Lipofectamine-3000, ThermoFisher) in serum-free medium (OptiMEM, Gibco). Transfection was performed for 48 hours, and cells were then collected by gentle flushing in PBS, enumerated and resuspended in commercial detection medium (HEK-Blue Detection, Invivogen), containing the substrate of the NFkB-reporter secreted embryonic alkaline phosphatase (SEAP). LPS stimulation was performed by incubating 50 000 transfected cells/well with 0.1-100 ng/mL LPS of *Escherichia coli* strain O111:B4 (LPS-EB, Invivogen) overnight in detection media in 96-wells plate (TPP). Activity of the NFkB reporter SEAP was measured by colorimetry at OD 655 nm.

### Recombinant protein expression and purification

A synthetic gene construct (Integrated DNA Technologies) for human FAM118B was cloned into a custom pET16 vector with an N-terminal 6xHis-hSUMO2 fusion tag, codon optimized for recombinant expression in *E. coli*, using Gibson assembly. N163A, H206A, and N163A/H206A double mutant FAM118B were all generated using ‘round the horn mutagenesis and the wild-type sequence as template. Sequencing results for plasmid preparations were confirmed before transformation into BL21-CodonPlus(DE3)-RIL *E. coli* (Agilent) for protein expression and subsequent purification as described below.

Transformant *E. coli* were plated on MDG-agar plates (0.5% glucose, 25 mM Na2HPO4, 25 mM KH2PO4, 50 mM NH4Cl, 5 mM Na2SO4, 2 mM MgSO4, 0.25% aspartic acid, 100 mg/mL ampicillin, 34 mg/mL chloramphenicol, and trace metals) and grown overnight at 37 °C. Single colonies were then inoculated into liquid MDG media and grown overnight at 37 °C, 230 rpm shaking. Overnight MDG cultures were used as inoculum for 1 L M9ZB minimal media liquid cultures (0.5% glycerol, 1% Cas-amino Acids, 47.8 mM Na2HPO4, 22 mM KH2PO4, 18.7 mM NH4Cl, 85.6 mM NaCl, 2 mM MgSO4, 100 mg/mL ampicillin, 34 mg/mL chloramphenicol, and trace metals). M9ZB cultures were grown at 37 °C, 230 rpm until OD600nm >2.5, rested on ice for 20 minutes and then induced with 0.5 mM IPTG and set to shake at 230 rpm overnight at 16 °C. Cells were pelleted by centrifugation, washed with PBS, and flash frozen in liquid N2 for storage at -80 °C.

Frozen cell pellets were resuspended in lysis buffer (20 mM HEPES-KOH pH 7.5, 400 mM NaCl, 10% glycerol, 30 mM imidazole, 1 mM DTT) and subjected to sonication to extract soluble protein. Cell lysates were clarified with centrifugation and purified using conventional gravity-flow Ni-NTA affinity chromatography. Briefly, lysate was loaded on Ni-NTA resin (Qiagen) equilibrated with lysis buffer and the flow-through was added back over the column once to increase chances of binding. Columns were then washed with lysis buffer supplemented to 1M NaCl, washed with lysis buffer, and protein was ultimately eluted in lysis buffer supplemented to 300 mM imidazole. Eluted protein was dialyzed overnight (20 mM HEPES-KOH pH 7.5, 250 mM KCl, 5% glycerol, 1 mM DTT) in the presence of human SENP2 protease (D364–L589, M497A) to remove the N-terminal 6xHis-SUMO2 solubility tag. Protein was concentrated post-dialysis using 30 kDa cutoff Amicon centrifuge filters and loaded onto a Superose-6 16/600 (Cytiva) gel filtration column (equilibrated in 20 mM HEPES-KOH pH 7.5, 250 mM KCl, 1 mM TCEP) connected to an NGC FPLC (Bio-Rad). Fractions were verified for protein content and purity via SDS-PAGE analysis and pooled fractions were further concentrated >10 mg/mL and flash frozen with liquid N2 for long-term storage at -80 °C.

### Fluorescence plate reader analysis of SIRanc enzymatic activity

50 μL reactions were built in PCR tubes before transfer to white-walled clear bottom 96-well plates. Reactions included FAM118B at 15 μM, nicotinamide 1,N(6)-ethenoadenine dinucleotide (ε-NAD) at 500 μM, 20 mM HEPES-KOH pH 7.5, 100 mM NaCl, and -/+ 10% PEG 400. All components were thawed on ice and reactions were incubated for 15 minutes at 37 °C prior to addition of fluorescent NAD^+^ analog and transfer to a Synergy H1 plate reader (BioTek). Fluorescence recordings at 300 nm excitation/410 nm emission were measured in kinetic mode every minute for one hour.

## Supplementary Material

### Supplementary Figures

**Figure S1.**
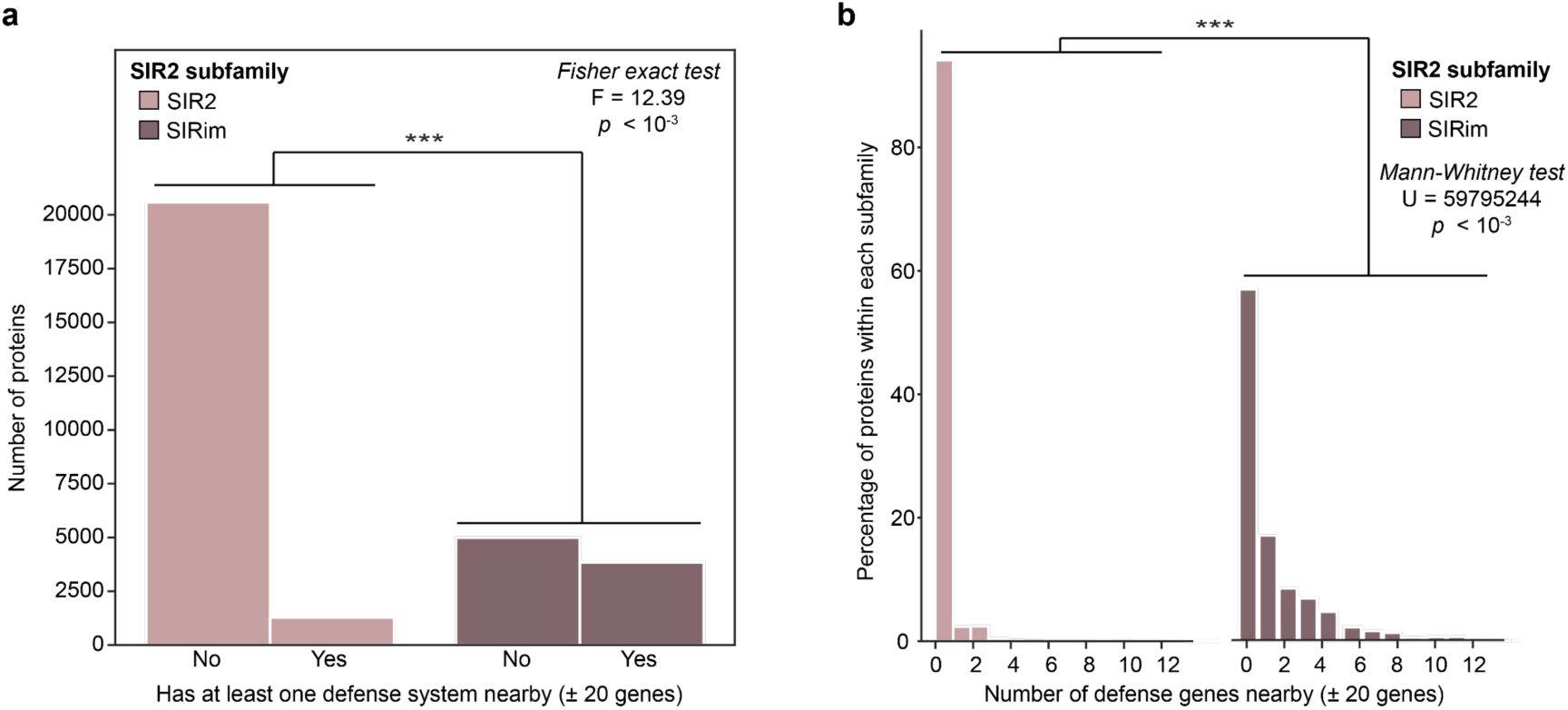
The genomic context around bacterial proteins from the SIRim subfamily is significantly enriched in known antiphage systems compared with the genomic context nearby bacterial SIR2 subfamily proteins. **a.** Number of bacterial proteins which have at least one defense system in their genomic neighborhood (20 genes upstream and 20 genes downstream). Proteins are grouped according to their SIR2 subfamily : SIR2 (beige) and SIRim (brown). Two sided Fisher exact test is used to assess statistical significance of the difference between the two distributions (***: *p*<10^-3^). **b.** Distribution of the number of defense genes in the genomic context of bacterial SIR2 family proteins per subfamily: SIR2 (beige) and SIRim (brown). The x-axis (“*Number of defense genes nearby*”) is duplicated across the two subfamilies for visualization purposes, but the y-axis (“*Percentage of proteins within each subfamily*”) is shared. Two-sided Mann-Whitney test is used to assess statistical significance of the difference between the two distributions (***: *p*<10^-3^).

**Figure S2.**
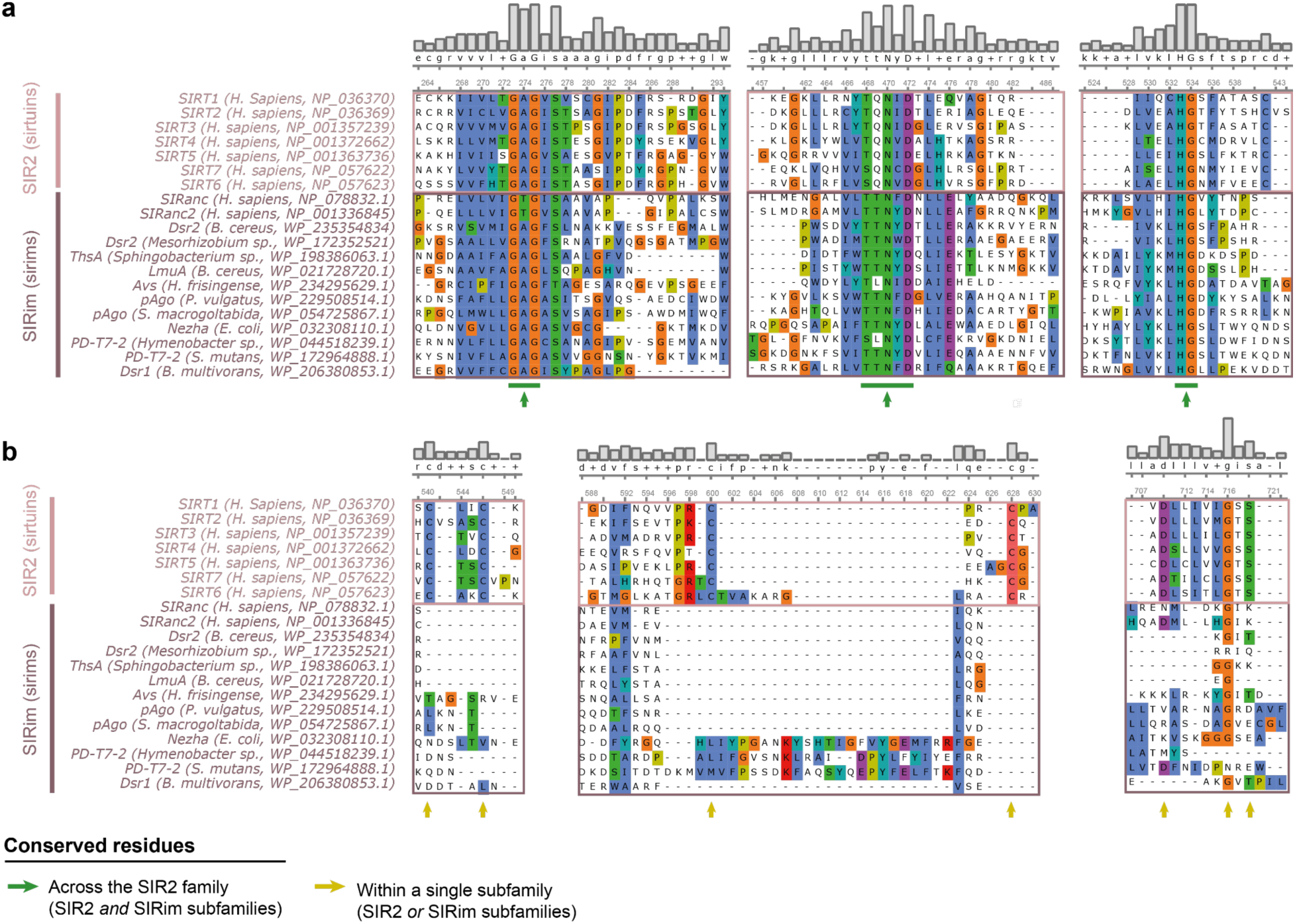
Multiple sequence alignment of twenty SIR2 family proteins sampled from bacteria and eukaryotes. Sequences are grouped by subfamily (SIR2: beige box, SIRim: brown box).Conserved residues are indicated by an arrow. **a.** Selected regions of the multiple sequence alignment which exhibit strong conservation both within subfamilies and among subfamilies. All the conserved motifs (G.G, ..N.D and .HG) are located within the NAD+ binding site of the proteins (*1–4*). **b.** Selected regions of the multiple sequence alignment which exhibit strong conservation within one of the two subfamilies only. The four cysteine (C) residues conserved within the SIR2 subfamily (beige) form a tetrad which coordinates a Zn^2+^ ion and forms the binding site of deacetylated amino acid residue (Fig 1b) (*1*). This tetrad is absent from all SIRim proteins (brown).

**Figure S3.**
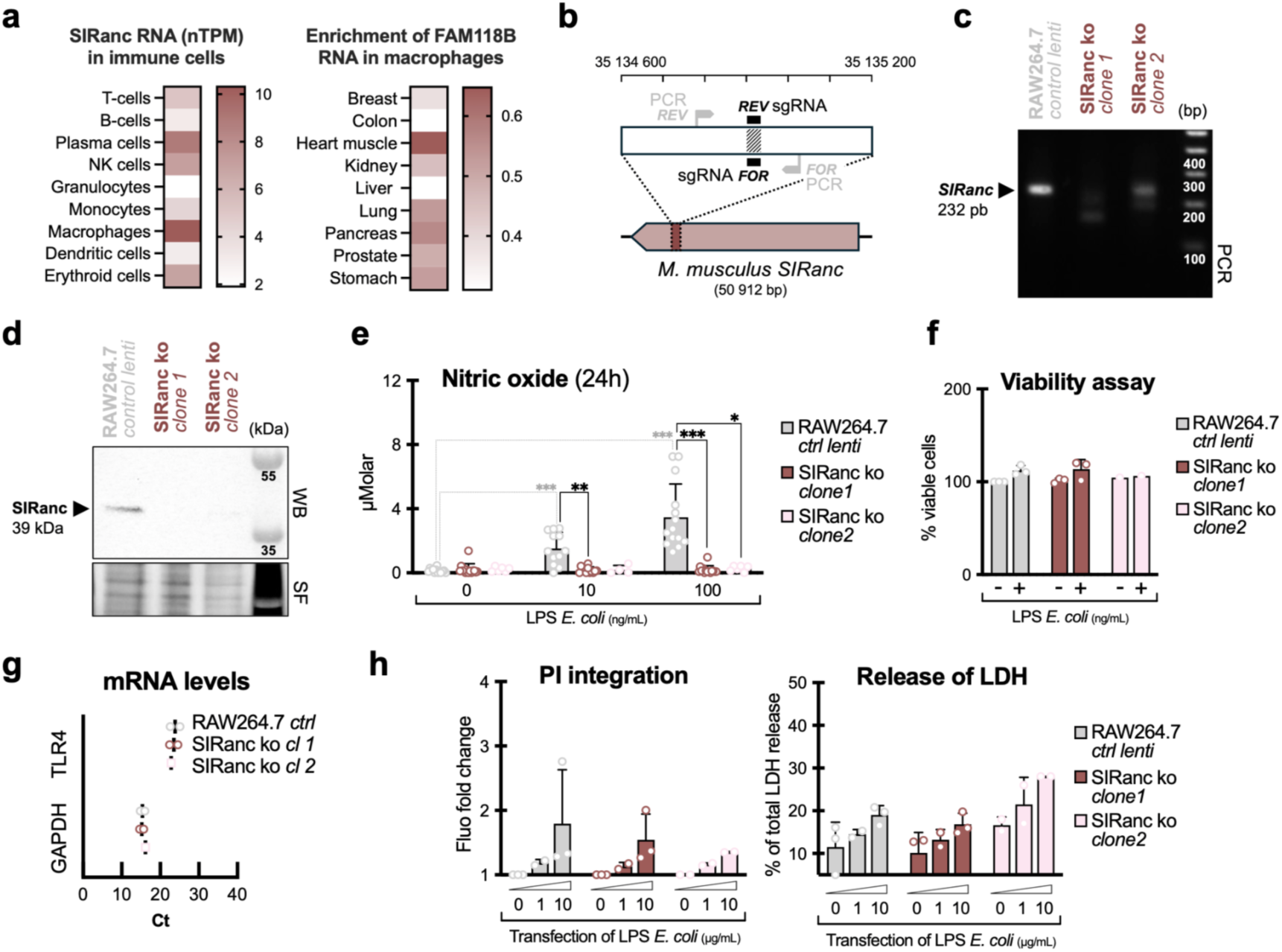
SIRanc knock-out (KO) in RAW264.7 murine macrophages. **a.** Heat maps of SIRanc RNA levels (number of transcripts per million, nTPM) in immune cells and enrichment ratio of SIRanc RNA in macrophages of various tissues. **b.** Knock-out strategy for SIRanc in RAW264.7 cells. **c.** PCR on genomic DNA of wild type or SIRanc KO (two clones) murine macrophages. **d.** Western blot & stain free (WB & SF) of wild type or SIRanc KO (two clones) murine macrophages. **e.** Wild type or SIRanc KO (two clones) murine macrophages were stimulated with LPS from *E. coli* to activate TLR4, and nitric oxide production was measured at 24h. **f.** Wild type or SIRanc KO (two clones) murine macrophages were stimulated with LPS from *E. coli* to activate TLR4 and viability was measured by Alamar Blue assay. **g.** RT-qPCR of GAPDH and TLR4 in wild type or SIRanc KO (two clones) murine macrophages. **h.** Propidium iodide integration and release of lactate dehydrogenase (LDH) in wild type or SIRanc KO (two clones) murine macrophages transfected overnight with 1-10 µg/mL of LPS from *E. coli*.

**Figure S4.**
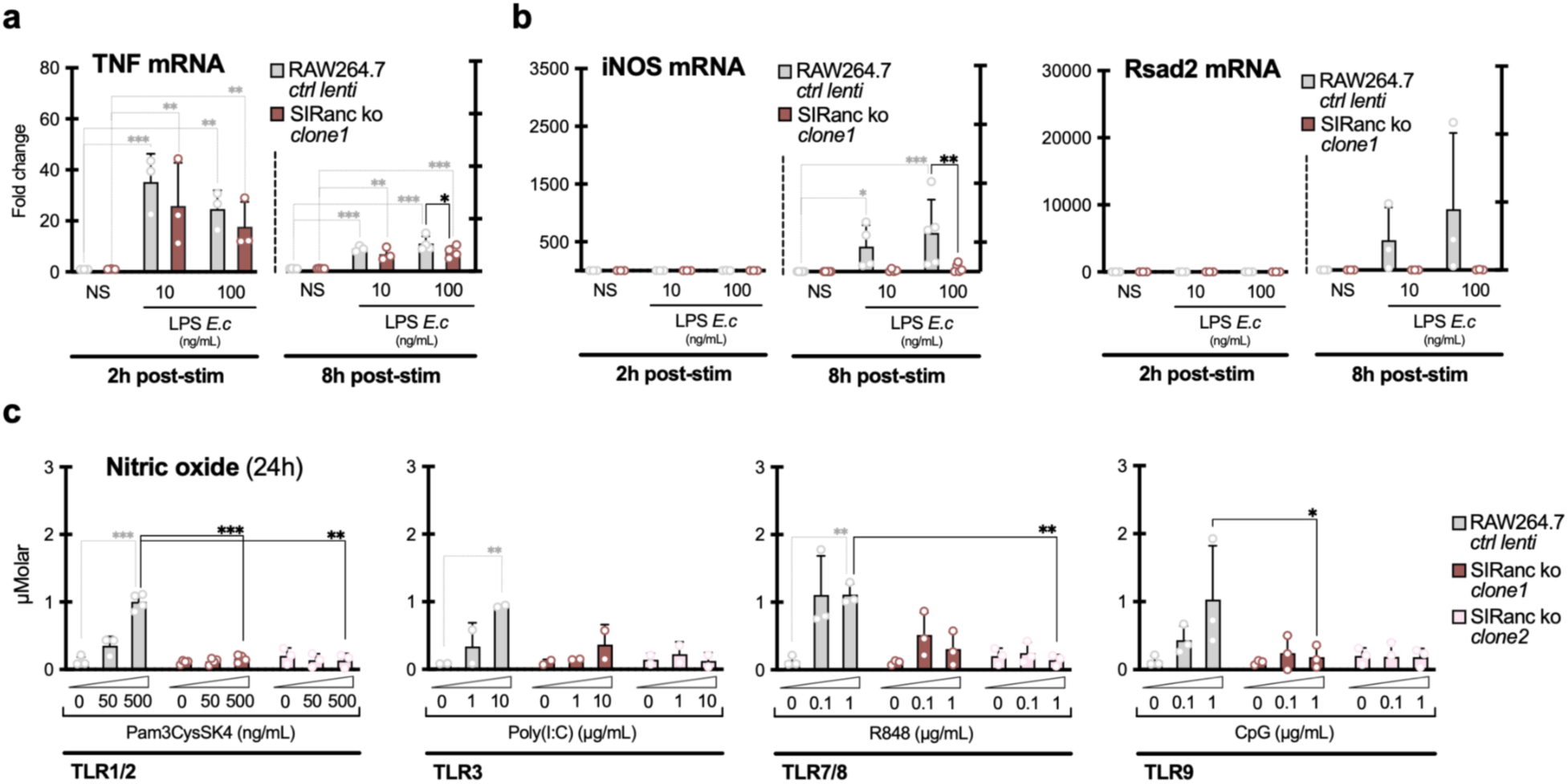
SIRanc contributes to TLRs signaling in RAW264.7 murine macrophages. **a-b**. Wild type or SIRanc KO murine macrophages were stimulated with LPS and levels of TNF, iNOS and Rsad2 transcripts were measured at 2h and 8h by RT-qPCR. **c**. Wild type or SIRanc KO (two clones) murine macrophages were stimulated with Pam3CysSK4, Poly(I:C), R848 or CpG, agonists of TLR1/2, TLR3, TLR7/8 and TLR9, respectively, and nitric oxide production was measured at 24h.

**Figure S5.**
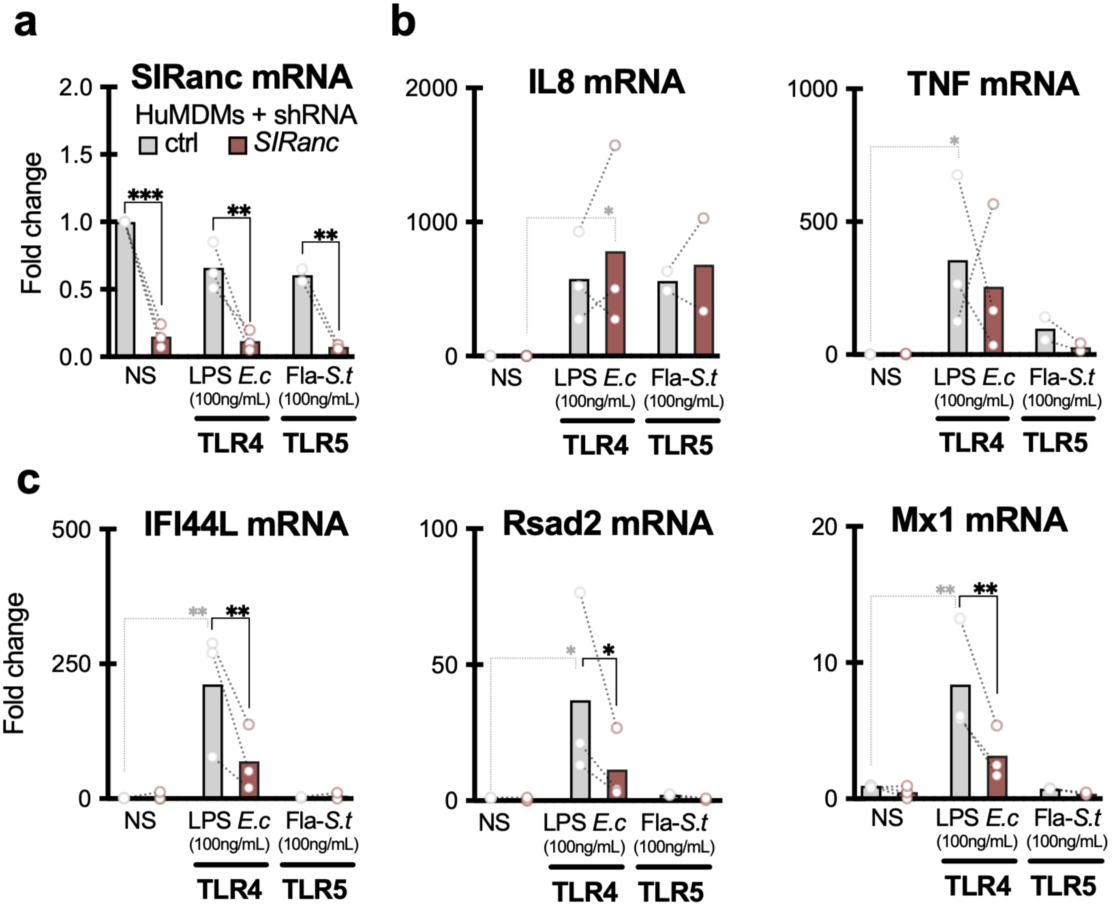
SIRanc contributes to TLR4 signaling in human MDMs. **a-c.** Human monocyte-derived macrophages stably expressing a control shRNA, or a shRNA downregulating the levels of SIRanc transcripts, were stimulated with LPS from *E. coli* or flagellin from *S. typhimurium*, to activate TLR4 and TLR5, respectively. Levels of SIRanc, IL8, TNF, IFI44L, Rsad2 and Mx1 transcripts were measured RT-qPCR at 8h post stimulation.

**Figure S6.**
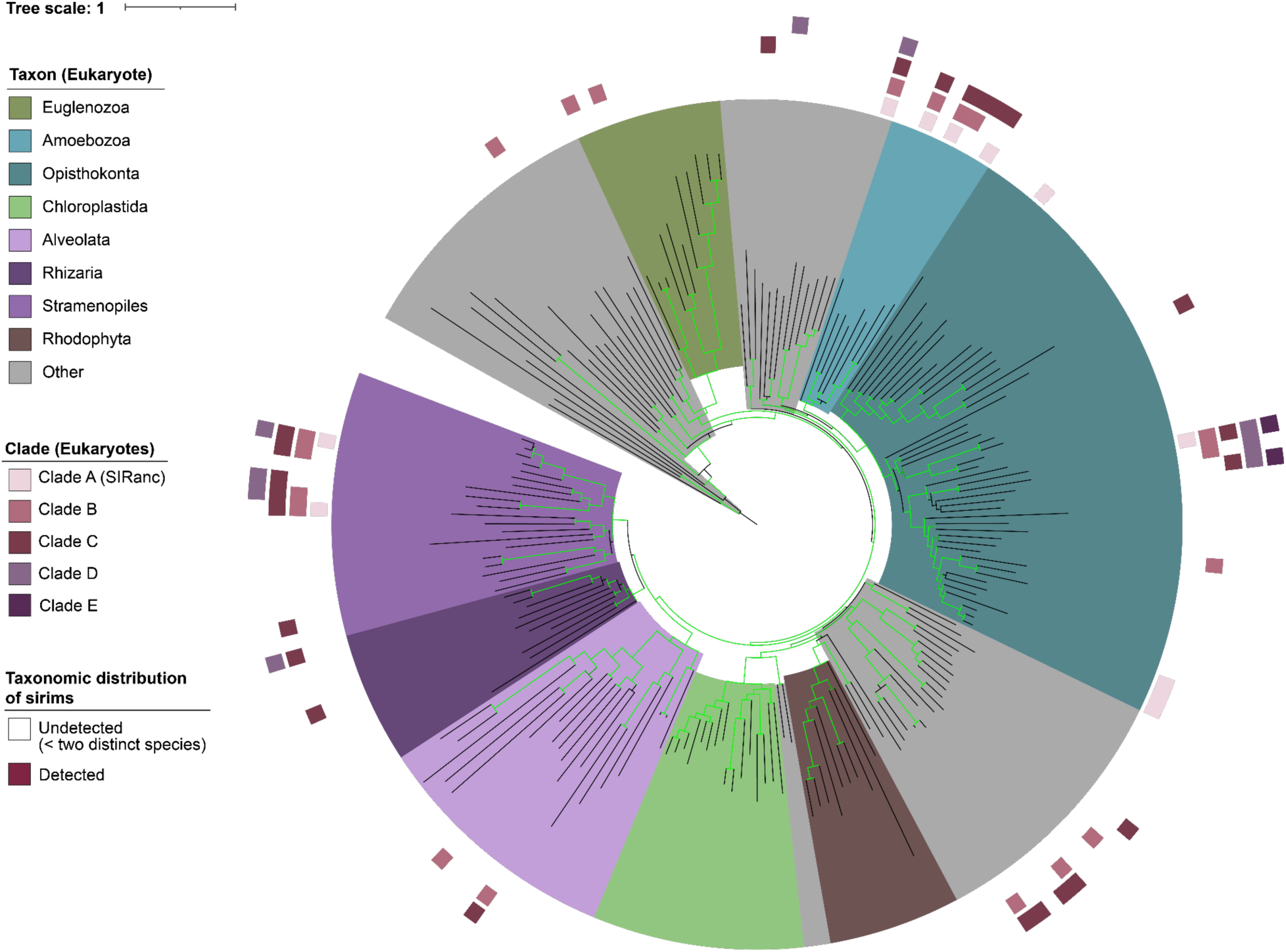
Distribution of the five major eukaryotic SIRim clades identified in eukaryotes. The phylogenetic tree from The Complete Set (TCS) of Eukprot (196 species) is used to draw the taxonomic distribution of the SIRim proteins identified in the Eukprot v3 database of eukaryotic genomes (985 species) (*5*). SIRim proteins found in species present in the Eukprot v3 database (985 species) but not in the TCS (196 species) are assigned to their closest relative present in the TCS tree. We consider that a sirim protein is detected in the taxon represented by a given leaf of the tree if (i) sirims are detected in at least two distinct species represented by this leaf or (ii) the leaf represents a single eukaryotic species in Eukprot v3 and sirims are detected within this species. Eukaryotic sirims are grouped into five distinct clades based on our phylogenetic analysis (Fig 4a, Fig S7 & S8). The tree is midpoint rooted. A pruned version of this tree can be found in Fig 4b.

**Figure S7.**
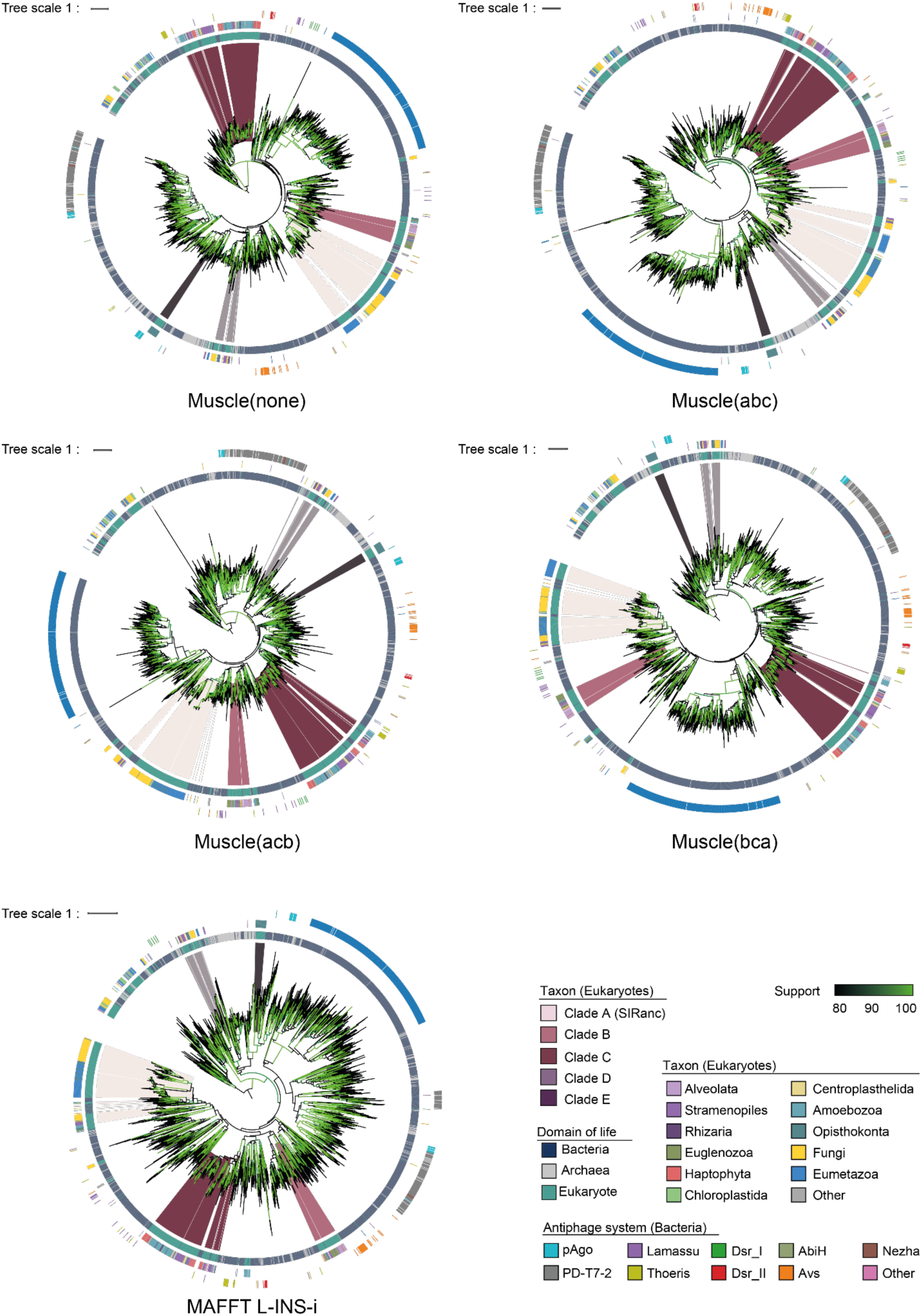
The five major clades of eukaryotic sirims are stably identified regardless of the sequence alignment method. The same representative sequences of bacterial, archaeal and eukaryotic sirims are used as in Fig 4a. Representative sequences are aligned on their SIRim domain using either Muscle super5 (first four trees) or MAFFT (last tree, bottom). The Multiple Sequence Alignment (MSA) generated using Muscle are also built by permuting the guide tree (permutations: “none”, “abc”, “acb”, “bca”). The domain of life from which each protein originates (inner ring, blue, gray, green), the taxon for eukaryotic sirims (middle ring) and the corresponding antiphage system type for bacterial sirims (outer ring) are also indicated on each tree. One phylogenetic tree is built from each MSA using IQTree. This procedure allows us to test the stability of the identified eukaryotic clades of sirims (clades A to E) to perturbations of the MSA used to build the phylogenetic tree. We observed that the five clades of eukaryotic sirims defined from the phylogenetic tree in Fig 4a are stably formed in all the obtained trees, indicating that they are significant. We also observe that the clades of eukaryotic sirims tend to associate with the same clades of bacterial antiphage sirims in all the trees.

**Figure S8.**
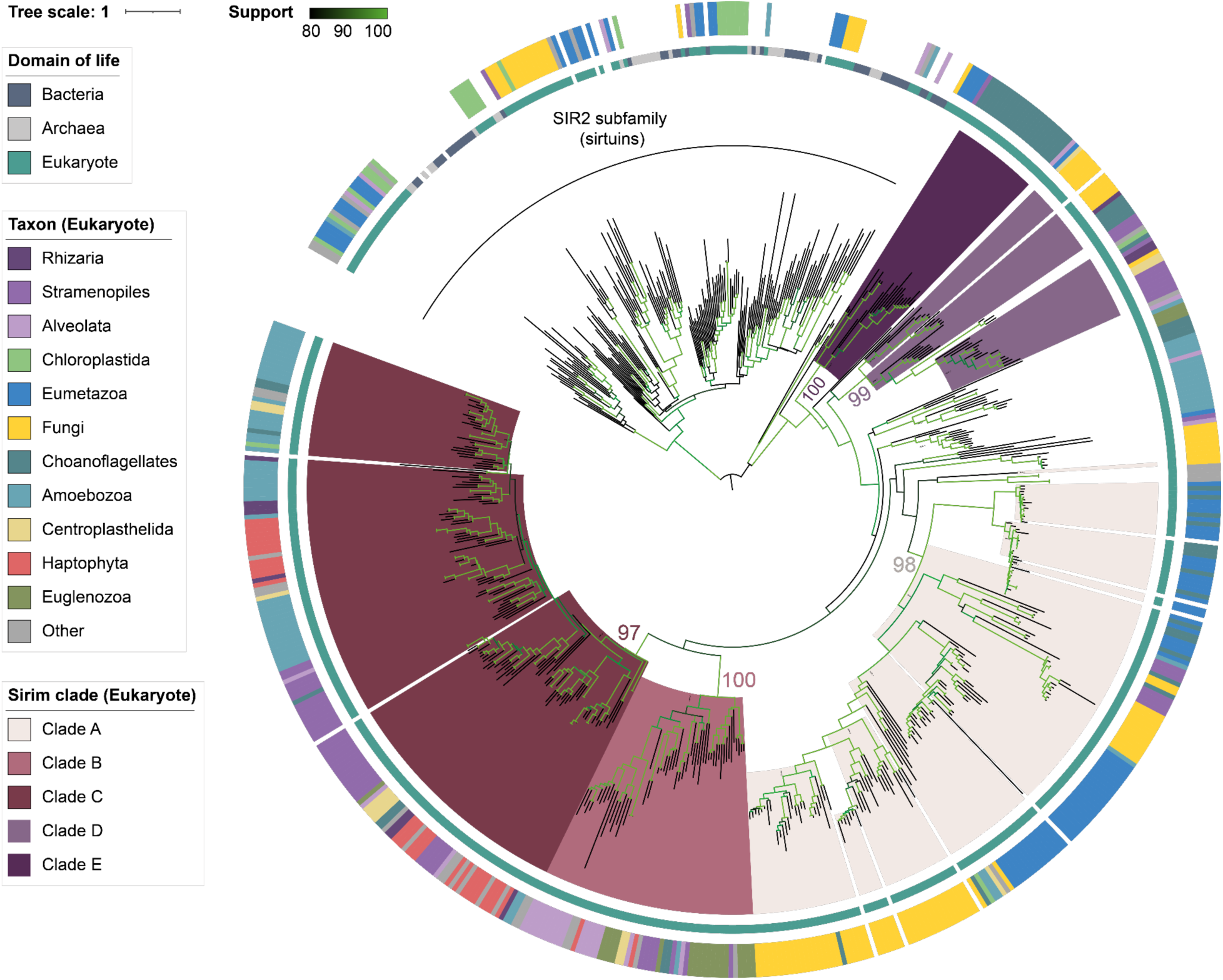
Phylogenetic tree of the eukaryotic SIRim. The tree only includes SIRim proteins from eukaryotic organisms and is rooted using the SIR2 subfamily of proteins (sirtuins) from across the tree of life as an outgroup. The five clades of eukaryotic sirims defined from the tree in Fig 4a are represented (inner ring) and are also retrieved in this tree with good support. Eukaryotic sirim clades left in white branch are unstable when rebuilding the trees, so are left apart from the classification of eukaryotic sirims. The domain of life (middle ring) and the eukaryotic taxon (outer ring) are also indicated. Shades of green on branches indicate UltraFast Bootstrap support values. Support values are explicitly provided for key branches defining the major eukaryotic sirim clades.

**Figure S9.**
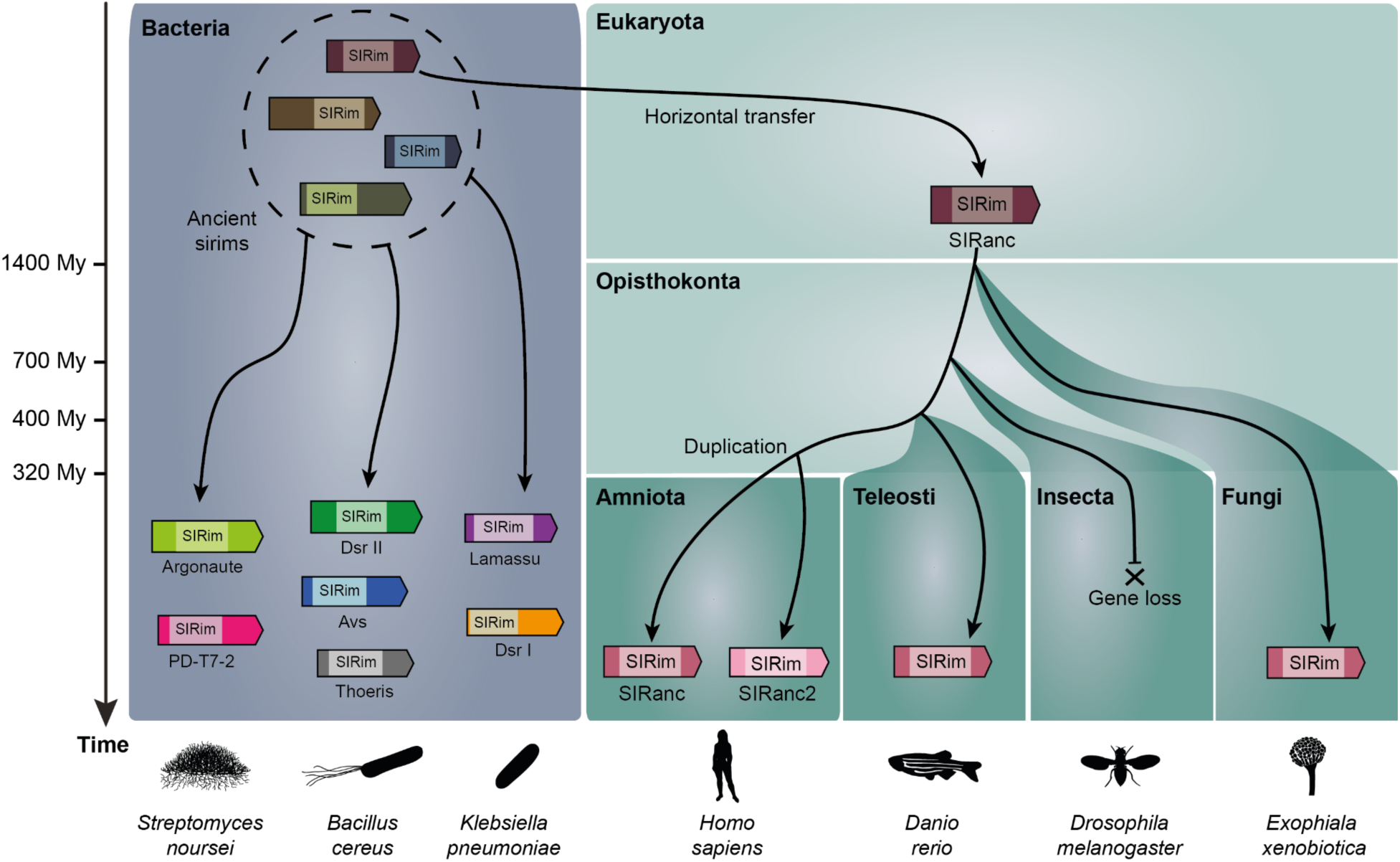
Proposed evolutionary scenario explaining the appearance of the SIRanc lineage through horizontal gene transfer from bacteria to eukaryotes and its subsequent diversification. The evolutionary scenario presented above exemplifies four major eukaryotic taxa: amniota, teleostei (bony fishes), insecta and fungi, even though SIRanc lineage proteins are found in other taxa. For each taxon, example organisms are provided at the bottom of the figure. The depicted time scale is arbitrary. In bacteria, the set of extant SIRim proteins present today originates from a pool of ancient SIRim proteins. In eukaryotes, homology detection and phylogenetic trees reveal several independent lineages of SIRim proteins, including the SIRanc lineage (Fig 4a, Fig S6-8, Clade A). This lineage is present in diverse eukaryotes including fungi and animals (opisthokonta), suggesting that it was present in eukaryotes prior the last common ancestor of opisthokonta more than 1,400 millions years ago (Fig 4a and Fig 4b, Fig S6-8, Clade A). Based on the trees, the ancestor to all SIRanc lineage proteins was probably acquired from an ancient bacterial sirim through horizontal gene transfer, and then diversified with the appearance of the diverse eukaryotic taxa. In some taxa of opisthokonts, the gene was lost (e.g. in insecta). In the rest of the taxa, the gene was maintained and can be found in current biological organisms. In the genome of amniota (a taxon that notably contains mammals and birds), two copies of the gene can be found (*SIRanc*, originally named *FAM118B*, and *SIRanc2*, named *FAM118A*). This suggests that the *SIRanc* gene underwent a duplication event that led to two paralogs. This gene duplication must have occurred prior to the last common ancestor of eukaryotes (more than 320 million years ago).

**Table S5.**
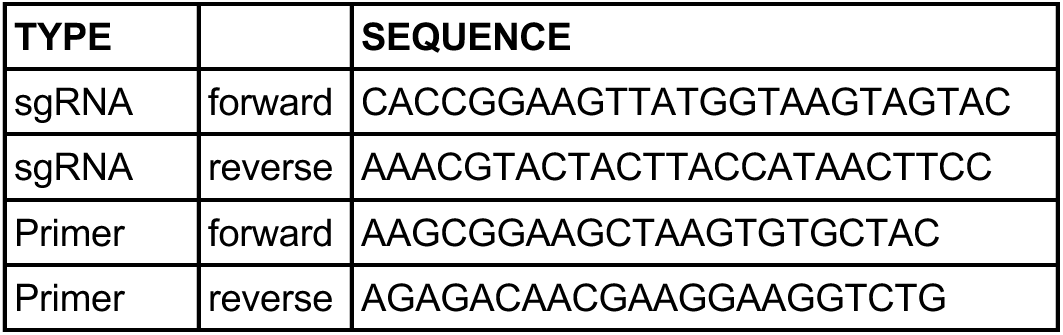
Primers and sgRNA used to construct RAW264.7 cells knocked-out for FAM118B.

**Table S6.** References of Taqman primers and probes used for RT-qPCR on RAW264.7 cells and human monocyte-derived macrophages.

| SPECIES | TARGET | REFERENCE |
| --- | --- | --- |
| Mouse | GAPDH | Mm99999915_g15 |
| Mouse | TNF | Mm00443258_m1 |
| Mouse | iNOS (Nos2) | Mm00440502_m1 |
| Mouse | Rsad2 | Mm00491265_m1 |
| Human | HPRT | Hs02800695_m1 |
| Human | FAM118B | Hs01090503_m1 |
| Human | IL-8 | Hs00174103_m1 |
| Human | IL-6 | Hs00174131_m1 |
| Human | TNF | Hs00174128_m1 |
| Human | IFI44L | Hs00915292_m1 |
| Human | Rsad2 | Hs00369813_m1 |
| Human | Mx1 | Hs00895608_m1 |

